# Evolutionary learning in the brain by heterosynaptic plasticity

**DOI:** 10.1101/2021.12.14.472260

**Authors:** Zedong Bi, Guozhang Chen, Dongping Yang, Yu Zhou, Liang Tian

## Abstract

How the brain modifies synapses to improve the performance of complicated networks remains one of the biggest mysteries in neuroscience. Canonical models suppose synaptic weights change according to pre- and post-synaptic activities (i.e., local plasticity rules), implementing gradient-descent algorithms. However, the lack of experimental evidence to confirm these models suggests that there may be important ingredients neglected by these models. For example, heterosynaptic plasticity, non-local rules mediated by inter-cellular signaling pathways, and the biological implementation of evolutionary algorithms (EA), another machine-learning paradigm that successfully trains large-scale neural networks, are seldom explored. Here we propose and systematically investigate an EA model of brain learning with non-local rules alone. Specifically, a population of agents are represented by different information routes in the brain, whose task performances are evaluated through gating on individual routes alternatively. The selection and reproduction of agents are realized by dopamine-guided heterosynaptic plasticity. Our EA model provides a framework to re-interpret the biological functions of dopamine, meta-plasticity of dendritic spines, memory replay, and the cooperative plasticity between the synapses within a dendritic neighborhood from a new and coherent aspect. Neural networks trained with the model exhibit analogous dynamics to the brain in cognitive tasks. Our EA model manifests broad competence to train spiking or analog neural networks with recurrent or feedforward architecture. Our EA model also demonstrates its powerful capability to train deep networks with biologically plausible binary weights in MNIST classification and Atari-game playing tasks with performance comparable with continuous-weight networks trained by gradient-based methods. Overall, our work leads to a fresh understanding of the brain learning mechanism unexplored by local rules and gradient-based algorithms.

## Introduction

Understanding the cellular-level learning mechanisms underlying brain intelligence is a grand aim of neuroscience. The outstanding question regarding this problem is how the brain adapts the synapses in its large-scale neuronal network to improve task performance: a question referred to as the ‘credit assignment problem’ [1].

Canonical models to solve the credit assignment problem suppose synaptic weights change according to pre- and post-synaptic activities (i.e., local plasticity rules, stemming from the Hebbian rule [2]), implementing gradient-descent algorithms. These models largely have two mainstreams [3]. (1) Backpropagation models. In these models, synaptic weights are updated according to locally available signals propagated through feedback connections [4, 5]. These signals can be either weight gradients [4, 5] or the targets of neuronal activities to be reached through weight updating [6, 7]. (2) Perturbation models. In these models, stochastic fluctuations are imposed on neuronal activities or synaptic weights [8, 9, 10]. The changes in task performance of a neural network due to these fluctuations are evaluated and broadcasted to all synapses through neuromodulators; weight gradients are then inferred from these fluctuated evaluations to guide synaptic updatings. However, despite some clues, experimental evidence to support these models remains controversial [3]. This inspires us to suspect that there may be important biological ingredients that current models neglect. Here we explore two such blind spots: one is an algorithmic paradigm other than gradient-based algorithms, and the other is a synaptic plasticity paradigm other than local rules, explained below:

Two algorithmic paradigms successfully train large-scale neural networks on real-world tasks in artificial intelligence. One is gradient-based algorithms, which inspire the gradient-based canonical algorithms discussed above; the other is evolutionary algorithms (EAs) [11], whose neural implementations are seldom discussed. In EA, a population of agents are evolving (**Figure 1a**). Each generation of agents are reproduced by the parental agents (i.e., the agents with higher fitness, or in other words, better task performance) in the previous generation through mutation or cross-over; elite agents (i.e., the agents with the highest fitness) become agents in the next generation without mutation, a technique called elitism. EA is a more general framework than gradient-descent algorithms: not only synaptic weights, but EA can also train network connections, which are fixed in gradient-descent algorithms [11]. It is unclear how EA may be implemented in the brain through biologically plausible mechanisms.

**Figure 1:**
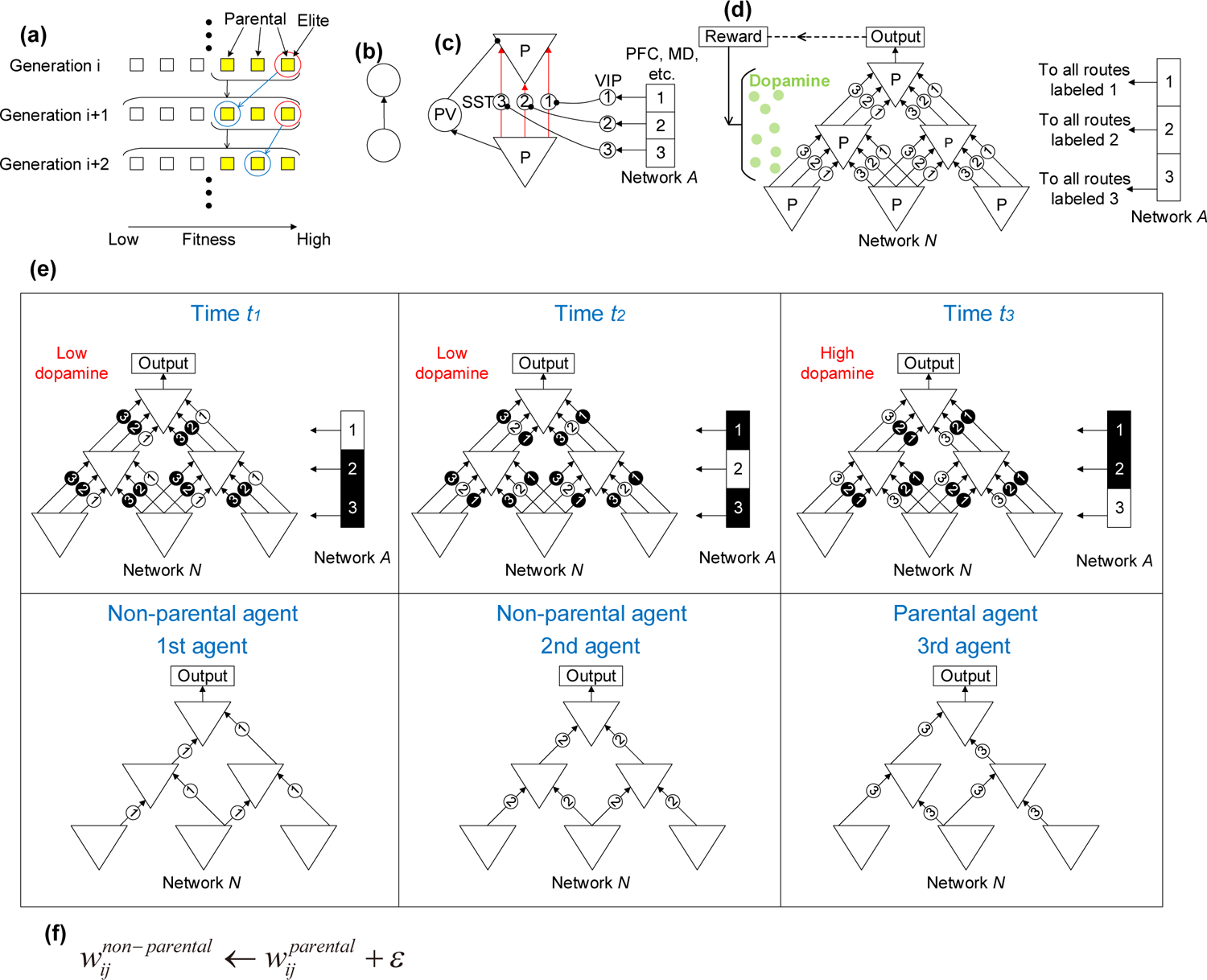
The network architecture. **(a)** Schematic of the evolutionary algorithm. Each generation of agents (squares) are reproduced by multiple parental agents (yellow squares) in the previous generation with high fitness (black arrows and brackets). The one with the highest fitness (red circle) is the elite and becomes an agent of the next generation without any mutation (blue arrows and circles), a technique called elitism. (**b**) Traditional models suppose that two neurons are connected by a single connection. (**c**) Our model supposes that two pyramidal neurons are connected through multiple dendritic routes, which are gated by MD or PFC (network *A*) through SST and VIP interneurons. Only synapses (red) between pyramidal neurons are plastic in our model, all the other synapses to and from PV, SST and VIP interneurons remain fixed. (**d**) Dendritic routes labeled *k* (*k* = 1, 2, 3) in network *N* are gated by neurons labeled *k* in network *A*. Dopamine, which represents the evaluation of the output of network *N*, are released dispersedly to network *N*. (**e**) Neurons in network *A* are activated alternately one at a time. At time *t_k_* (*k* = 1, 2, 3), only neuron *k* in network *A* is activated (white square) with others silent (black squares), so that the dendritic routes labeled *k* in network *N* are gated on (white circles) with others gated off (black circles). The sub-network connected by all the *k*th dendritic routes in network *N* is regarded as the *k*th agent in evolutionary algorithm (lower subplots). The 3rd agent stimulates a high level of dopamine (red text in the upper-right subplot), is therefore the parental agent; the other two agents stimulate a low level of dopamine, are therefore non-parental agents. (**f**) The plasticity rule. The synaptic efficacy *w*^non-parental^ from the *j*th to *i*th pyramidal neurons in a non-parental agent becomes the synaptic efficacy between the same pair of pyramidal neurons in a parental agent plus a small value *ɛ*.

Most brain learning mechanisms are proposed based on the assumption of locality, i.e., the change of a synapse depends on its pre- and post-synaptic activities, stemming from the Hebbian rule [2]. However, synaptic plasticity is not entirely local: a synapse can be changed by the activities of neighboring neurons unconnected with the synapse through inter-cellular signaling pathways, a phenomenon called heterosynaptic plasticity [12, 13]. For example, the diffusive nitric oxide (NO) associated with a synapse potentiated by pre- and post-synaptic spike pairing can potentiate other neighboring synapses [14, 15]; the pre-synaptic release of glutamate can stimulate long-range propagating calcium waves in astrocytes, depressing synapses in a broader spatial range [16, 17]. Blockade of heterosynaptic plasticity significantly impairs cognition [18]. It has been proposed that heterosynaptic plasticity is important for synaptic homeostasis during Hebbian learning [12], whose role, though necessary, is auxiliary. It remains unexplored whether heterosynaptic plasticity alone may imply a powerful learning paradigm.

Here we propose an EA model that is biologically realized by heterosynaptic plasticity. In our model, the brain contains many different information routes, each representing an EA agent. The performances of these agents are evaluated through alternatively gating on each route with the others gated off. Agent selection and reproduction are realized by dopamine-guided heterosynaptic plasticity. To examine heterosynaptic plasticity alone, we do not add any local rule in the model.

We show that our EA model achieves three significant merits simultaneously:

1. Biological consistency. Our EA model is based on a broad spectrum of cellular-level experimental evidence, including heterosynaptic potentiation mediated by NO [14, 15], heterosynaptic depression mediated by astrocyte calcium wave [16, 17], dendritic gating [19], binary synapses [20], dopamine gating of synaptic plasticity [21], meta-plasticity [22, 23], memory replay [24], cooperative plasticity between the synapses within a dendritic neighborhood [25, 26, 27], etc. With these biological elements, our EA model trains recurrent neural networks to exhibit dynamics analogous to brain dynamics in experimental observations [28].
2. Broad competence. Our EA model can well train neural networks with either feedforward or recurrent network architecture made of either analog or spiking neurons, which maximizes the generalizability of the learning mechanism.
3. Powerful capability. Our EA model can train deep networks with biologically plausible binary weights in MNIST classification and Atari-game playing tasks up to performance comparable with continuous-weight networks trained by gradient-based algorithms.

Collectively, our work sheds light on a new perspective on understanding the learning mechanism underlying brain intelligence.

## Results

### Network architecture

In traditional computational models of neuroscience, two neurons are connected by a single connection (**Figure 1b**). This contrasts with the reality that in the brain two pyramidal neurons are connected by synapses on multiple dendrites (**Figure 1c**) [29]. These dendritic routes can be independently gated by dendrite-inhibiting somatostatin (SST) interneurons [19], which are controlled by a hub area, e.g., prefrontal cortex (PFC) or mediodorsal thalamus (MD) [30], through long-distance corticocortical or thalamocortical connections to vasoactive intestinal peptide (VIP) interneurons that inhibit SST interneurons [31, 32]. Parvalbumin (PV) interneurons inhibit pyramidal neurons near the soma [19].

This neural substance provides a natural basis for implementing EA (**Figure 1a**), illustrated in **Figure 1d, e**, where we suppose that the dendrites of pyramidal neurons in a network **N** representing hippocampus or neocortex are gated by a network *A* representing PFC or MD. If neuron *k* in network *A* is active, all the dendrites labeled *k* in network *N* are gated on, with the others gated off. Neurons in network *A* are alternatively activated one at a time, such that dendrites with different labels are gated-on alternatively (**Figure 1e, upper subplots**). With different dendrites gated on, the network *N* gives different outputs in response to the same stimulus, leading to different rewards, either from the real world or estimated by the brain [33]. These reward signals stimulate different levels of dopamine, informed dispersedly to the whole network *N* to guide synaptic plasticity [21, 34] (**Figure 1d**). The sub-network of the network *N* connected by the dendrites labeled *k* is regarded as the *k*th agent in EA. Dopamine level stimulated by an agent indicates the fitness of that agents, such that high (or low) dopamine level indicates parental (or non-parental) agents (**Figure 1e, lower subplots**).

In the next subsection, we will show that the heterosynaptic plasticity between pyramidal neurons is equivalent to a learning rule in which the synapse *w*^non-parental^ from the *j*th to *i*th pyramidal neurons on a non-parental dendrite becomes approximately the synapse *w*^parental^ on a parental dendrite (**Figure 1f**). In this way, non-parental agents are updated to be descendants of the parental agent with some mutations (represented by *ϵ* in **Figure 1f**) caused by the stochastic nature of the nervous system [35, 36]. All the synapses to and from inhibitory interneurons are kept fixed in our model (**Figure 1c**). Note that we only consider mutation in this EA, and do not consider cross-over, which however can easily be included for further research (see Discussion).

### Basic learning process

We consider a delayed reward process widely in nature or the laboratory: neuronal activities result in eligibility traces in synapses, according to which sparse and delayed rewards then guide synaptic plasticity. Eligibility traces are molecular states in synapses, which can change synaptic efficacies under a high dopamine level, but have no effect under a low dopamine level [37]. The above plasticity rule (**Figure 1f**) can be realized by heterosynaptic plasticity through the following cellular-level mechanisms. To illustrate the mechanisms, we consider two pre-synaptic neurons collected to one post-synaptic neuron through three dendrites (*a*, *b*, *c*) with binary synapse efficacies [20] (represented by large or small boutons in **Figure 2**); at a certain time step, dendrite *c* is gated on (see **Figure 2**).

1. If a synapse on the gated-on dendrite *c* has large efficacy (e.g., synapse *c*_1_ from neuron 1 in **Figure 2a**), a pre-synaptic spike from neuron 1 will be very likely to stimulate a post-synaptic spike. This pairing of pre- and post-synaptic spikes produces diffusive messengers such as nitric oxide (NO) [38, 14]. This NO and simultaneously arriving spikes from neuron 1 generate *high* potentiation eligibility traces in both *c*_1_ and neighboring synapses (*a*_1_ and *b*_1_) from the same pre-synaptic neuron (i.e., neuron 1) on gated-off dendrites *a* and *b* [15, 14, 39], see **Figure 2c**. This heterosynaptic potentiation is subject to a spatial scope comparable with the scale of the dendritic arbor of a pyramidal neuron (∼ 150 *µ*m in hippocampal CA1 [15, 14]). Synapses from neuron 2 (i.e., synapses *a*_2_, *b*_2_ and *c*_2_) cannot be potentiated by the diffusive NO without simultaneous pre-synaptic spikes.
2. If a synapse on a gated-on dendrite *c* has small efficacy (e.g., synapse *c*_2_ from neuron 2 in **Figure 2b**), a pre-synaptic spike from neuron 2 will be unlikely to stimulate a post-synaptic spike, resulting in *low* potentiation eligibility traces in both *c*_2_ and neighboring synapses (*a*_2_ and *b*_2_) from the same pre-synaptic neuron (i.e., neuron 2) on gated-off dendrites *a* and *b*, see **Figure 2c**.
3. Pre-synaptic release of glutamate stimulates astrocytes to spread calcium waves [40, 17] (**Figure 2a, b**), which induces depression eligibility traces in synapses spatially widespread [41, 42, 16] (at least 300∼500 *µ*m in hippocampal CA1 [43, 16]). The depression eligibility traces on different synapses are supposed to be similar in our model due to the widespread spatial scale, see **Figure 2c**.

**Figure 2:**
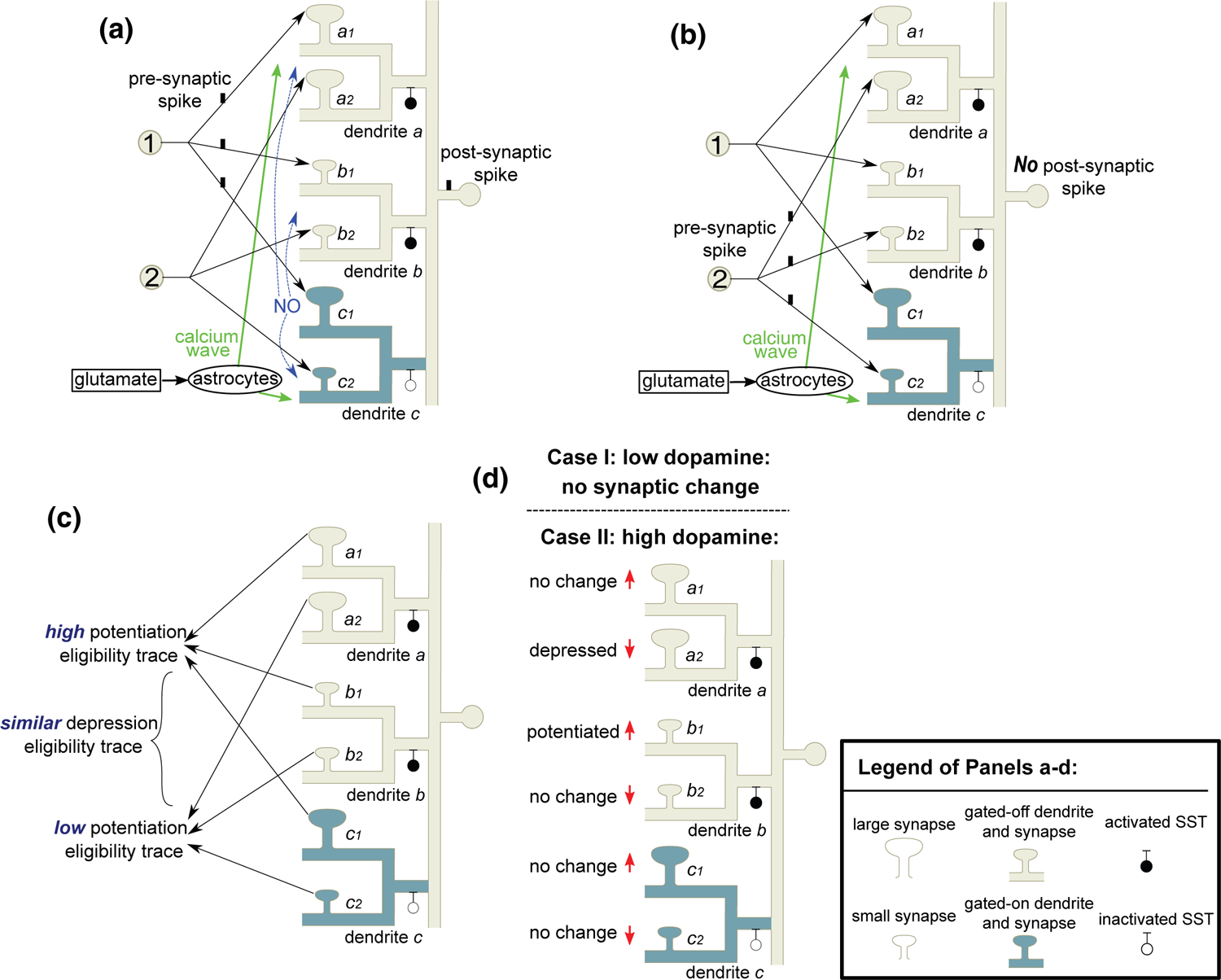
Heterosynaptic plasticity induces EA. (**a**) When neuron 1 emits a spike, the large synapse *c*_1_ on the gated-on dendrite *c* stimulates a post-synaptic spike; this pairing of pre- and post-synaptic spikes produces diffusive NO, which, together with the spikes simultaneously invading synapses *a*_1_ and *b*_1_ from neuron 1, induces potentiation eligibility traces in these synapses. (**b**) When neuron 2 emits a spike, the small synapse *c*_2_ on the gated-on dendrite *c* does not stimulate a post-synaptic spike, therefore no potentiation eligibility traces in synapses *a*_2_ and *b*_2_. In both panels **a** and **b**, pre-synaptic release of glutamate stimulates astrocytes to produce widely spreading calcium waves, which induce depression eligibility traces in all synapses. (**c**) The synapses targeted by neuron 1 have higher potentiation eligibility traces than those targeted by neuron 2, but the depression eligibility traces in these synapses are similar. (**d**) Case I: with a low dopamine level, no synaptic change happens. Case II: with a high dopamine level, the synapses *a*_1_, *b*_1_, *c*_1_ (or *a*_2_, *b*_2_, *c*_2_) with high (or low) potentiation eligibility traces tend to be potentiated (or depressed), represented by upper (or lower) red arrows. Synapses *a*_1_, *c*_1_ (or *b*_2_, *c*_2_) remain unchanged because they already have large (or small) efficacy.

Collectively, if a synapse on a gated-on dendrite has high (or low) efficacy, it will induce high (or low) potentiation eligibility traces in the neighboring synapses from the same pre-synaptic neuron on gated-off dendrites; the depression eligibility traces on different synapses are similar (**Figure 2c**). The mechanisms above are best established in the hippocampus [12] (also see the references above), and have also been observed in the neocortex [13, 39, 42, 44, 45].

If the delayed reward is small so that the dopamine level is low (i.e., the gated-on dendrite *c* is non-parental, see **Figure 1e**), no synaptic change will happen (Case I in **Figure 2d**). If the delayed reward is large so that the dopamine level is high (i.e., the gated-on dendrite *c* is parental, see **Figure 1e**), the synapses will be updated [21] according to the following rules (Case II in **Figure 2d**): synapses with large potentiation eligibility traces will be potentiated, while synapses with small potentiation eligibility traces will be depressed by the depression eligibility trace. As biological synapses take binary efficacies [20], potentiation (or depression) only works in synapses with low (or high) efficacy, whereas those already with high (or low) efficacy remain unchanged (**Figure 2d**). As a result, synapses on non-parental dendrites are updated towards those on parental dendrites connecting the same pair of pre- and post-synaptic neurons (i.e., synapses *a*_1_ and *b*_1_ are updated towards *c*_1_, whereas *a*_2_ and *b*_2_ towards *c*_2_), while those on parental dendrites (i.e., synapses *c*_1_ and *c*_2_) remain unchanged (**Figure 2d**), fulfilling the requirement of the EA (**Figure 1f**).

Generally speaking, the above arguments still hold if we regard each neuron in the model as a population of closely neighboring pyramidal neurons, so that the synapses *a*_1_, *b*_1_, *c*_1_ (or *a*_2_, *b*_2_, *c*_2_) do not have the same pre- or post-synaptic neurons. On one hand, both the heterosynaptic potentiation and depression mechanisms (**Figure 2a, b**) are mediated by inter-cellular pathways, therefore still hold even if the synapses belong to different pre- or post-synaptic neurons, as long as these synapses are spatially nearby [39, 15, 16]. On the other hand, closely neighboring pyramidal neurons have high firing synchrony [46], so the simultaneity of NO and spike required to potentiate *b*_1_ and *c*_1_ in **Figure 2a** can be realized if the spikes invading *a*_1_, *b*_1_ and *c*_1_ are not from the same pre-synaptic neuron, but from different neighboring pre-synaptic neurons with high firing synchrony. We will discuss more about synchronous pre-synaptic firing below (**Figure 5**).

### Multiple parents and elitism realized by meta-plasticity

In EA of computer science, each generation usually has multiple parents and at least one elite [47, 48] (**Figure 1a**). Here we show that both multiple parents and elitism can be implemented in our model by introducing a meta-plasticity mechanism. In this mechanism, if a synapse with large (or small) efficacy on a gated-on dendrite receives potentiation (or depression) signals under high dopamine level, the efficacy of this synapse will get stabler at the large (or small) level, harder to be switched in future learning processes (**Figure 3a**).

**Figure 3:**
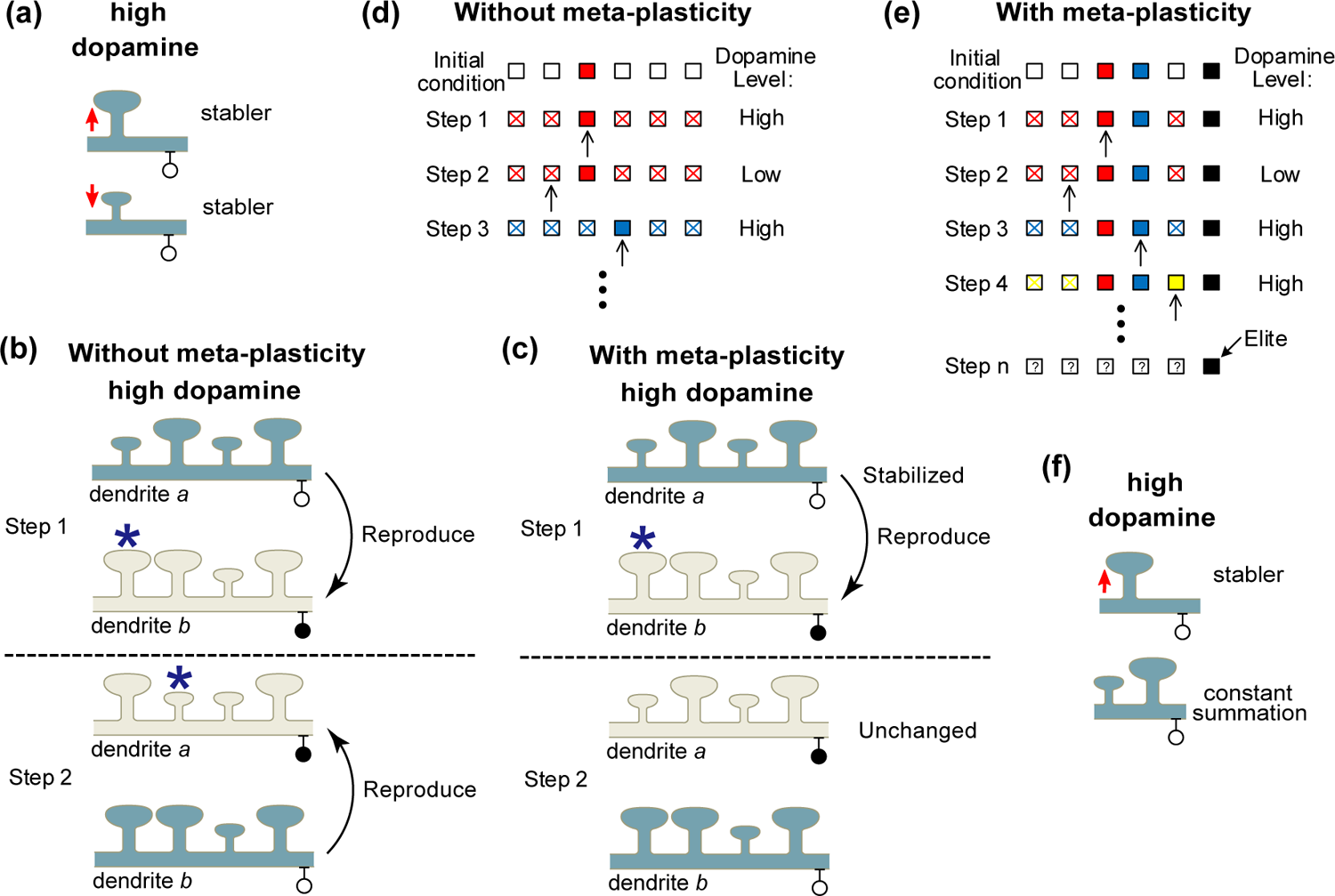
Multiple parents and elitism realized by meta-plasticity. (**a**) Schematic of the meta-plasticity. On a gated-on dendrite (white circles represent inactivated SST interneurons, see the legend of **Figure 2**), if a large (upper panel) or small (lower panel) synapse receives potentiation or depression signal (upward or downward arrow) at a high dopamine level, the synapse will be stabilized at large or small efficacy. (**b**) Schematic of the case without meta-plasticity. At time step 1 (or 2), the gated-on dendrite *a* (or *b*) is parental, so a high dopamine level is induced at both time steps. At step 1 (upper panel), the synaptic configuration of dendrite *a* is reproduced to dendrite *b*. At step 2 (lower panel), the synaptic configuration of dendrite *b* is reproduced to dendrite *a*. The asterisks indicate mutations (i.e., *ɛ* in **Figure 1f**) that happen with a small probability. (**c**) With meta-plasticity, the synapses on dendrite *a* are stabilized when dendrite *a* is parental (upper panel), so that when dendrite *b* is parental at a later time step (lower panel), the synapses on dendrite *a* will not change. (**d**) The learning process without meta-plasticity. Each box represents an agent in EA, i.e., a dendritic route in **Figure 1e**. The initial condition contains one parental agent (red square) that can induce a high dopamine level and five non-parental agents (white squares). In step 1, the red parental agents are activated (black arrow), so that all non-parental agents are updated towards the red agent, becoming its descendants (red crosses). In step 2, the activated agent cannot induce a high dopamine level, so this agent remains non-parental. In step 3, the activated agent induces a high dopamine level, so all the other agents (including the red parental agent) are updated toward the activated agent, becoming its descendants (blue crosses). We see that the agents at every time step have a single parent and no elite without meta-plasticity. (**e**) The learning process with meta-plasticity. Notice that the parental agents (red, blue, yellow and black squares) are stabilized, so that in step 1, 3 or 4, non-activated parental agents (i.e., the parental agents not indicated by the black arrow) do not update toward the activated parental agent (i.e., the parental agent indicated by the black arrow) that stimulates high dopamine level. After many steps (last row), parental agents may also lose their synaptic configurations (the question marks mean we do not know the states of those agents); but the elite agent (black square), which can induce the highest dopamine level, is the stablest one, and can last for the longest time. (**f**) The stabilization of small synapses can be realized by stabilizing large synapses (upper panel) together with dendrite-level synaptic homeostasis (lower panel).

To understand the effect of this meta-plasticity, suppose dendrite *a* is parental at time step 1. Without the meta-plasticity, if another dendrite *b* is parental at a later time step 2, synapses on dendrite *b* will reproduce their efficacies on dendrite *a* (**Figure 3b, lower panel**), under the mechanism illustrated in Case II of **Figure 2d**. In this case, dendrite *b* will become the parent of dendrite *a*. With the meta-plasticity, however, the synapses on dendrite *a* are stabilized, without changing toward the synapses on dendrite *b*, so that dendrite *b* will not become the parent of dendrite *a* (**Figure 3c, lower panel**). After considering the training process of multiple dendrites, we find that without the meta-plasticity, all the dendrites at any step have a single parent (**Figure 3d**); with the meta-plasticity, however, the dendrites can have multiple parents (**Figure 3e**). Besides, even with the meta-plasticity, synapses on a stabilized dendrite may also gradually lose their synaptic configuration after a sufficiently long time. The parental dendrite that induces the highest dopamine level (i.e., the elite) has the highest stability, and therefore will be maintained for the longest time, fulfilling the elitism technique in EA (**Figure 3e, black boxes**).

Physiologically, it is well known that dopamine stabilizes potentiated synapses of pyramidal neurons, after the excitability of pyramidal neurons is increased by the inhibition of SST or PV interneurons [22, 49, 50] (upper subplot of **Figure 3a**). However, the stablization of small synapses (lower subplot of **Figure 3a**), despite some clues [51, 52], remains to be experimentally tested. A better-established process to fulfill the stabilization of small synapses is dendrite-level synaptic homeostasis [53, 54], which constrains the total synaptic efficacy on a dendrite (**Figure 3f**). Under this constraint, small synapses are hard to be potentiated if other large synapses on the same dendrite are stabilized, realizing the stabilization of small synapses. More generally, if we regard a single neuron in **Figure 2** as a population of closely neighboring biological pyramidal neurons (as we discussed in the last paragraph of the previous subsection), and the information route is gated at the neuron level (instead of the dendrite level), neuron-level synaptic homeostasis [55, 56, 57] will fulfill the stabilization task.

### EA is compatible with off-line replay

According to the no-free-lunch theorem [58], no single learning algorithm is advantageous in every problem. Therefore, it is necessary for the brain to combine a variety of learning strategies for the best living of the animal, and it is also necessary for us to understand the suitable scenarios for EA, thereby getting insight into the functional roles of EA. Here we show that EA is compatiable with off-line replay.

Consider the following on-line learning scenario which possibly happens in a waking animal (Scenario 1, **Figure 4a**). In a learning session, the performance of a gated-on dendritic route in response to an input stimulus is scored by the reward feedback from the real world, inducing a corresponding dopamine level; learning sessions may be separated by intervals of variable durations. This scenario is not compatible with EA. To see this, consider a situation where dendritic route 1 brings 10 scores of reward in response to stimulus 1, and brings 0 scores in response to stimulus 2; dendritic route 2 brings 6 scores of reward in response to both stimuli 1 and 2 (**Figure 4c**). If high dopamine is induced only in the session corresponding to the highest reward (i.e., the session with stimulus 1 and dendritic route 1 gated-on), dendritic route 1 will be parental, and the synapses on dendritic route 2 will update towards those on dendritic route 1. However, this parental selection is incorrect because it is dendritic route 2 that brings more rewards on average over the two input stimuli (**Figure 4c**).

**Figure 4:**
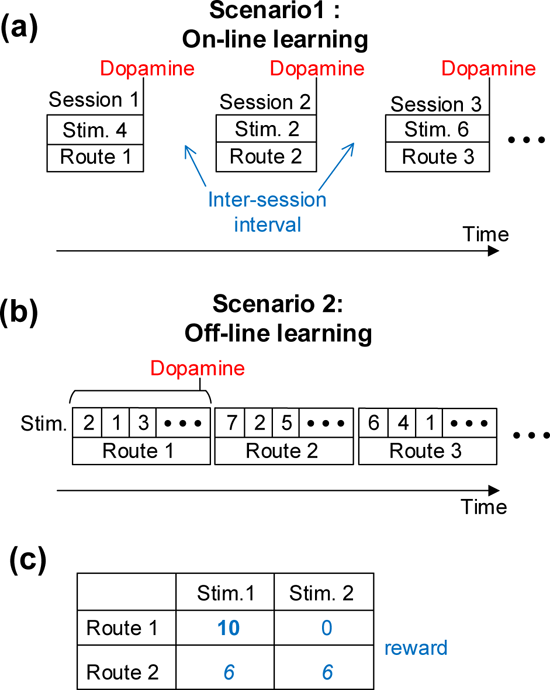
EA is compatible with off-line replay. (**a**) During an on-line learning scenario, the performance of a gated-on dendritic route in response to an input stimulus is represented by the level of dopamine given at the end of each session. There are intervals of variable durations between sessions (blue text). (**b**) During an off-line learning scenario, the brain can quickly sweep over samples of input stimuli when a dendritic route *k* (*k* = 1, 2, 3, · · ·) is gated on. The dopamine level represents the average reward associated with the replayed stimuli when the same dendritic route is gated on. (**c**) An example situation. Route 1 brings the highest reward (bold blue text) in response to a single stimulus, but route 2 brings the highest average reward (italic blue text) over both stimuli.

This problem can be solved if we suppose that the evaluation of dendritic routes happens during off-line replay (Scenario 2, **Figure 4b**): a gated-on dendritic route is quickly evaluated over a number of replayed inputs, and the dopamine level is released according to the average reward (estimated by the brain) associated with a number of input stimuli just replayed (**Figure 4b**). In this case, the dopamine level indicates the average performance of the gated-on dendritic route over a batch of input samples just replayed. If we set the batch size to be 2, the dendritic route 2 will be correctly selected as the parental agent during the replay in the case of **Figure 4c**.

The incompatibility of EA with on-line learning lies in the very nature of EA. In gradient-descent algorithms, the updating direction of a synaptic weight is computable and equal to the average over the directions computed from individual input stimuli samples. Therefore, the neural network can be optimized by accumulating small updating steps in response to individual stimuli during the on-line learning in Scenario 1. However, in EA, such synaptic updating direction can not be computed, and relies on the comparison of different dendritic routes. This comparison depends on the performance of dendritic routes over a number of stimuli samples (as an estimation of the performance over the whole stimuli set) instead of a single stimulus, which makes the on-line learning inappropriate. The compatibility of EA with off-line replay suggests that EA plays an important role in transforming episodic memory into unconscious dexterous skills at rest or in sleep [59, 60]. This off-line replay scenario (**Figure 4b**) is a prediction of our model, which can be experimentally tested in the future. In the following simulations, EA is performed in an off-line manner.

### Cooperative plasticity between synapses from synchronous inputs

From **Figure 2a**, the simultaneous arrival of the pre-synaptic spikes from neuron 1 at synapses *a*_1_, *b*_1_ and *c*_1_ is a key factor for the potentiation of *a*_1_ and *b*_1_. The inter-cellular diffusion of NO suggests that such potentiation still happens even if the spikes invading *a*_1_ and *b*_1_ do not come from neuron 1, but from a different neuron with synchronous firing with neuron 1. This subsection will investigate the EA process when single post-synaptic neurons receive from synchronous inputs from different pre-synaptic neurons.

Pre-synaptic neurons with synchronous activities target close dendritic locations [61, 62] through synapses with cooperative plasticity: when a synapse is potentiated, the neighboring synapses can also be potentiated at the same time [27] or become easy to be potentiated by only weak stimulation [25]; when a synapse is depressed, the neighboring synapses can also be depressed at the same time [26]. In our model, the synapses from synchronous-firing pre-synaptic neurons to the same post-synaptic dendrite are mutated downward or upward simultaneously (group mutation, GM) instead of independently (individual mutation, IM), modeling the cooperative plasticity between neighboring synapses due to cellular mechanisms in the post-synaptic dendrite [25, 26]. We found that GM is essential for the success of EA learning.

To illustrate the mechanisms, we consider three pre-synaptic neurons (1, 2, 3) with synchronous activities targeting a post-synaptic neuron through two dendrites (*a*, *b*), see **Figure 5a, b**. At a certain time, dendrite *a* is gated on while dendrite *b* is gated off, and the synapses from the three pre-synaptic neurons on dendrite *a* have large efficacies at the beginning (**Figure 5a, left column**). Under IM, each synapse is mutated independently with low probability. In this case, suppose synapse *a*_3_ on dendrite *a* is mutated downward (**Figure 5a, upper row**). High NO is emitted from the large-efficacy synapses *a*_1_ and *a*_2_ on the gated-on dendrite *a* (**Figure 5a, upper row, middle column**) when the spikes from neurons 1 and 2 invading *a*_1_ and *a*_2_. When this happens, the spikes from neurons 1 and 2 arrive at synapses *b*_1_ and *b*_2_ simultaneously, and the spikes from neuron 3 also arrive at synapses *a*_3_ and *b*_3_ simultaneously due to the synchronous activity of neuron 3 with neurons 1 and 2. Therefore, there are high potentiation eligibility traces in all the synapses (*a*_1_, *a*_2_, *a*_3_, *b*_1_, *b*_2_, *b*_3_) on both dendrites (**Figure 5a, upper row, middle column**). If the gated-on dendrite *a* is parental so that high dopamine is induced, all these synapses will become large (**Figure 5a, upper row, right column**): this means that the parental synaptic configuration on the parental dendrite *a* after IM (i.e., synapses *a*_1_, *a*_2_ and *a*_3_ have large, large and small efficacies respectively) cannot be reproduced to dendrite *b*, which contradicts with the EA requirement (**Figure 1f**). Under GM, however, the synapses from the three pre-synaptic neurons on dendrite *a* are mutated downward simultaneously (**Figure 5a, lower row, middle column**). In this case, the NO received by the synapses on both dendrites is low, inducing low potentiation eligibility traces. If dendrite *a* is parental so that high dopamine is induced, all the synapses in both dendrites will become small (**Figure 5a, lower row, right column**): this means that the parental synaptic configuration on the parental dendrite *a* after GM (i.e., synapses *a*_1_, *a*_2_ and *a*_3_ all have small efficacies) gets reproduced to dendrite *b*. A similar situation happens when the three synapses on dendrite *a* have small efficacies at the beginning (**Figure 5b, left column**): if dendrite *a* is parental (i.e., induces high dopamine) after IM (or GM), its mutated configuration will not (or will) be reproduced to dendrite *b* (**Figure 5b**), inconsistent (or consistent) with the EA requirement (**Figure 1f**).

**Figure 5:**
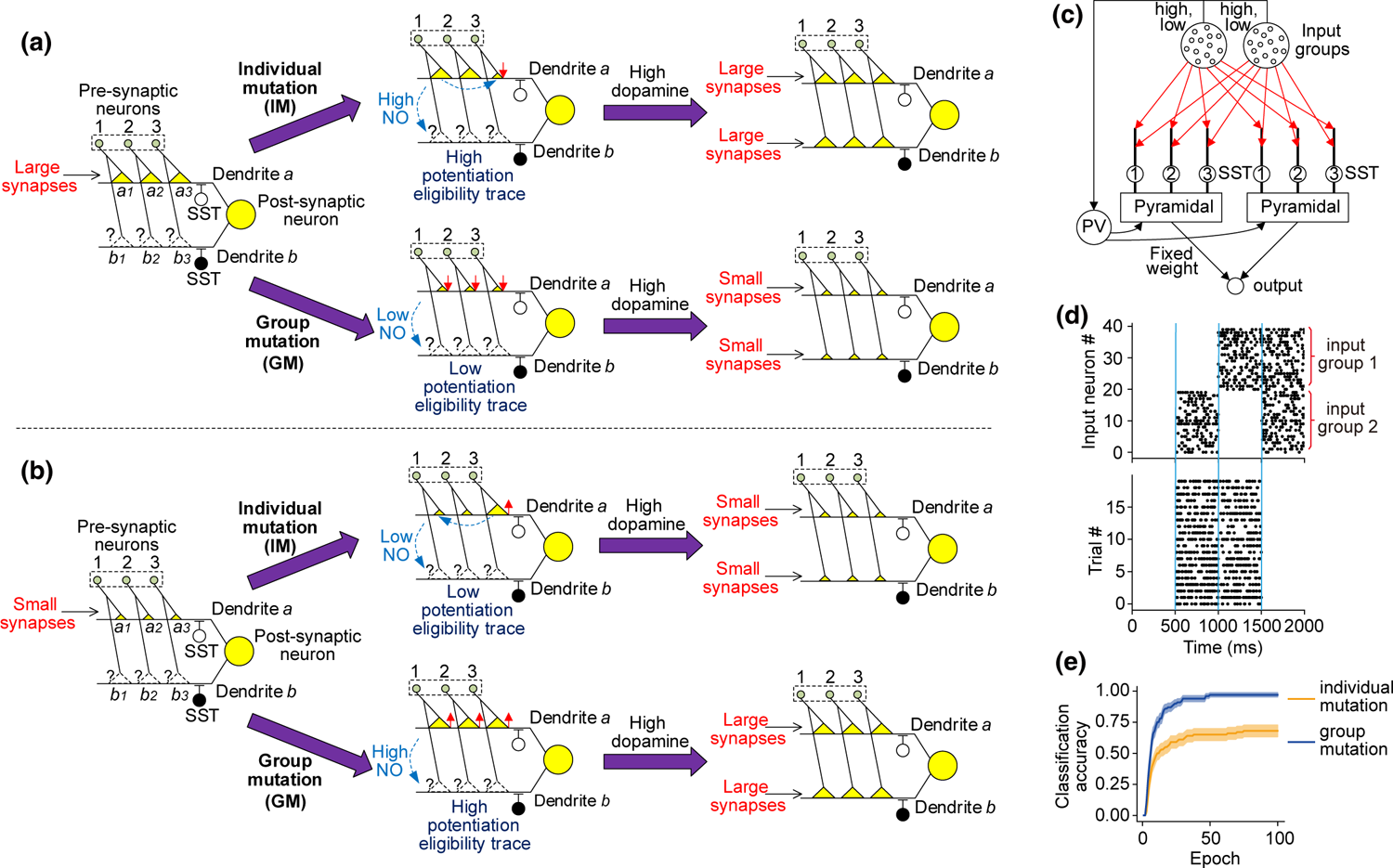
The computational function of the cooperative plasticity. (**a, b**) Schematic to illustrate the advantage of group mutation. The green circles in dashed boxes represent pre-synaptic neurons with synchronous activity. Large (or small) yellow triangles represent large (or small) synapses. Dashed triangles with question marks represent synapses with no matter large or small efficacy. Dendrite *a* or *b* is gated on or off by inactivated or activated SST interneurons (white or black circle). Upward (or downward) arrows indicate that the synapses are mutated upward (or downward). If the gated-on dendrite *a* is parental (i.e., induces high dopamine) after IM (or GM), its mutated configuration will not (or will) be reproduced to dendrite *b*. (**c**) The structure of the network to perform the XOR task. Pyramidal neurons receive from two input groups (whose activity levels can be high or low) through synapses (red arrows) on dendrites gated by SST interneurons. Red synapses are to be changed during training, and the other synapses are fixed. (**d**) Raster plot of the input neurons in one simulation trial (upper panel) and the single output neuron in multiple trials (lower panel) after training under group mutation. Vertical blue lines delimit four intervals in which the two input groups have different high or low activities. (**e**) Classification accuracy during the training process for individual (orange) or group (blue) mutation. Error belts indicate the standard error of the mean (s.e.m.) over 100 training trials.

We demonstrate the above mechanism by training a spiking neural network on the XOR task. In this network (**Figure 5c**), a population of pyramidal neurons receives Poisson spike trains emitted from two neuronal groups, and then give output to a single neuron. The synapses from the two input groups to the pyramidal neurons (i.e., red arrows in **Figure 5c**) should be trained so that the output neuron has a high firing rate when one input group has a high firing rate while the other is silent, and the output neuron has a low firing rate when both input groups have high or low firing rates (**Figure 5d**). In this model, neurons belonging to the same input group have correlated activities (**Figure 5d, upper panel**), and therefore have higher firing synchrony than the neurons belonging two different input groups.

We examined IM, when individual synapses are mutated independently, and GM, when the synapses from different neurons in the same input group to the same post-synaptic dendrite are mutated simultaneously. GM is a better mutation strategy than IM, because GM enables the network to quickly achieve high classification accuracy, whereas IM saturates the network at a low performance (**Figure 5e**). Therefore, cooperative plasticity between synapses from synchronous input neurons is necessary for EA to train the brain’s neuronal network successfully. See more details in **Supplementary Figure 1**.

### EA-trained neural networks mimic brain dynamics

To further illustrate the biological plausibility of EA, we trained recurrent neural networks to perform a context-dependent decision-making task [28] by EA, and compared the dynamics of EA-trained neural networks with that of the brain and backpropagation (BP)-trained neural networks [28]. In this task, neural networks are to make a binary choice according to one of its two input channels (representing the motion or color coherence of the random dots in the experiment of [28]) depending on whether the context signal is motion or color (**Figure 6a**). In the motion (color) context, the network should choose 1 or 2 depending on whether the motion (color) coherence is positive or negative, neglecting the color (motion) signal. The neural networks contain firing-rate units, each representing a group of spiking neurons (e.g., an input group in **Figure 5c**) with correlated activities in the brain. We trained the neural networks using the following simple algorithm (*EA-Simple*) to model the group mutation (**Figure 5a, b**) of the synapses from the same neuronal group:

1. Each synaptic weight takes binary values, one positive Δ*w*, one negative −Δ*w*, modeling the effective synaptic weight when the excitatory synapse has large or small efficacy under the global inhibition of PV interneurons (**Figure 5c**).
2. Descendants are mutated from parental agents with a small probability of synaptic flip. In other words, *ɛ* (**Figure 1f**) is usually zero, but with a small probability 2Δ*w* (or −2Δ*w*) if *w*^parental^ is −Δ*w* (or Δ*w*).

**Figure 6:**
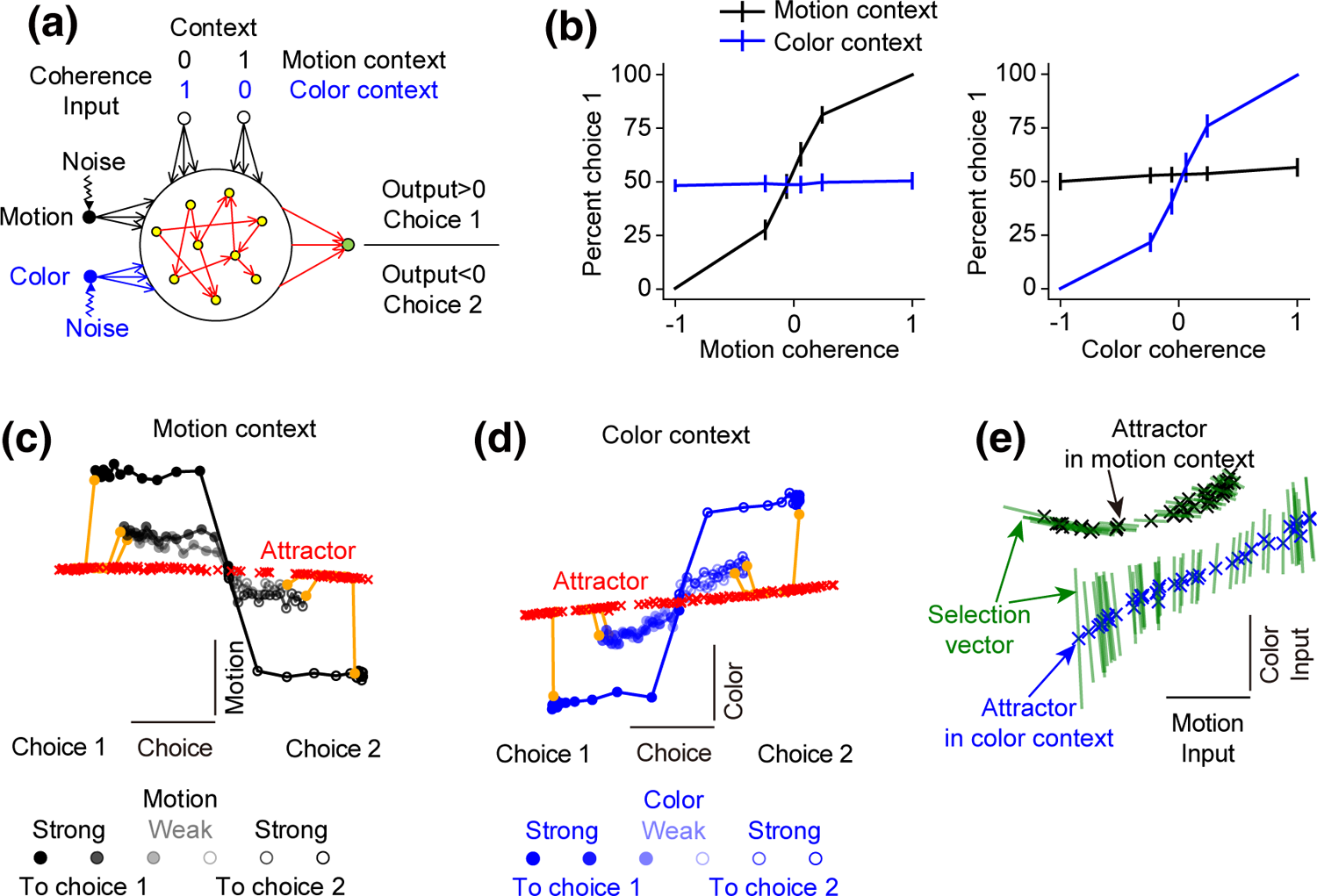
EA-trained neural networks mimic brain dynamics. (**a**) The architecture of the neural network model. In motion (or color) context, the neural network is to choose 1 or 2 depending on whether the input signal that indicates the motion (or color) coherence is larger than zero. (**b**) Psychometric curves show the percentage of choice 1 as a function of the motion and color coherence in the motion (black) and color (blue) contexts. Error bars represent s.e.m. over 8 training trials. (**c**) Dynamics of model population responses in the motion context in the subspace spanned by the axes of choice and motion, when the motion coherence takes different values. Each fixed point (red cross) has an eigenvalue close to zero, these fixed points together approximate a line attractor. After the inputs are turned off (orange dots and lines), the responses relax back towards the line attractor. Analogous to Figures 2 and 5 of [28]. (**d**) Similar to panel **c**, but in the color context and in the subspace spanned by the axes of choice and color. (**e**) The line attractor (black or blue crosses) and selection vector (green) at each fixed point, in the subspace spanned by the motion and color input weights. Analogous to Figure 6c of [28].

EA-trained networks mimic the brain and BP-trained networks in the following four aspects [28]. First, EA-trained networks successfully learned to perform the task, which is reflected in the behavioral psychometric functions (**Figure 6b**). Second, when making different choices, the trajectories of the network states move parallelly to a line attractor toward different directions at a distance proportional to the coherence strength (**Figure 6c, d**). Third, after defining three axes that respectively capture the most across-trial variance in the state space due to the choice, the motion and color coherence, we found that context usually has no substantial effect on the directions of the axes of choice, motion and color (**Supplementary Figure 2c**). Fourth, the selection vectors (i.e., the left eigenvector of the largest eigenvalue) of a line of attractors are aligned with the motion (or color) input and orthogonal to the color (or motion) input in the motion (or color) context (**Figure 6e**), which indicates that the relevant input pushes the network state along the direction of the line attractor, whereas the irrelevant input has no effect. These results imply that EA-trained networks mimic the dynamics of the brain in cognitive tasks, which further supports the biological plausibility of EA. See more details in **Supplementary Figure 2**.

### EA is a broadly competent and powerful learning paradigm

To demonstrate that EA is competent for various network architectures and capable of training neural networks for complicated tasks, we trained neural networks with biologically plausible binary weights using the EA-Simple algorithm on various tasks.

We first trained a sparse recurrent spiking neural network (SNN) to output arbitrary trajectory with time [63, 64] (**Figure 7a**, see Methods). Despite the binary weights of the recurrent and output synapses, the network’s output can well approximate the target trajectory (**Figure 7b, Supplementary Figure 3a**).

**Figure 7:**
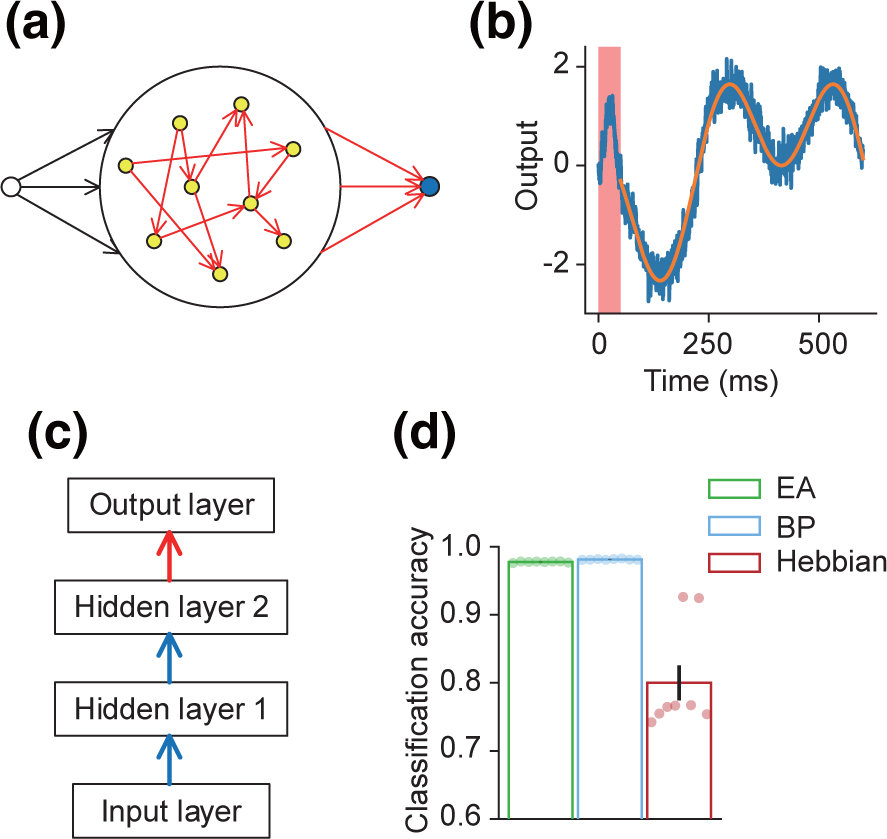
Training spiking and analog neural networks with recurrent and feedforward architectures using EA. (**a**) Schematic of the SNN to generate trajectories. Recurrent units (yellow circles) are sparsely connected. The output unit (blue circle) is trained to generate an arbitrary trajectory. Both the recurrent and output connections (red arrows) are binary and plastic. (**b**) An example of the actual output (blue) and target (orange) trajectory. The input signal is on during the initial 50 ms (red shading) and off afterward. (**c**) The architecture of the ANN to classify MNIST images. When studying the Hebbian algorithm in panel **d**, the lower layers were trained using the Hebbian algorithm (blue arrows), and the top layer was trained by gradient-descent algorithm supervisedly (red arrows). (**d**) Classification accuracy on the test dataset when training the deep network using BP, EA and Hebbian rule. Error bars represent s.e.m. over 8 training trials.

We then trained a feedforward analog neural network (ANN) with two hidden layers (**Figure 7c**) to classify the MNIST images. The final classification performance is comparable with continuous-weight networks with the same structure trained by BP (**Figure 7d, Supplementary Figure 3b**). We also trained continuous-weight networks layer-by-layer using a competitive Hebbian algorithm [65], except that the last layer was trained by gradient-descent algorithm supervisedly (**Figure 7c**), and found worse final performance (**Figure 7d, third bar**). Interestingly, if we supervisedly trained the last layer on top of the first hidden layer, the performance got better than the two-hidden-layer case (**Supplementary Figure 3c, d**), similar to the finding in [66]. This phenomenon implies that Hebbian algorithms cannot coordinate weights in different layers to fulfill better performance.

We further demonstrate the capability of the EA by training deep neural networks with binary weights to play Atari games, whose performance is comparable to continuous-weight networks trained by gradient-based methods such as DQN and A3C [67, 68] and EA-based method [69] (**Table 1**). We also tried a Hebbian-Q algorithm, where low-level layers were trained by a competitive Hebbian algorithm [65], and the top layer was trained by the Q learning [67], and found that the resulted performance was significantly worse than DQN, A3C and EA (**Table 1**). These results demonstrate that the biologically supported EA is a broadly competent algorithm capable of training deep neural networks on complicated tasks.

**Table 1:** Binary-weight networks trained by EA are competitive with continuous-weight networks trained by DQN and A3C in Atari games. In the fifth column, bold black numbers indicate the scores of EA-trained binary-weight networks that are higher than both the scores of DQN- and A3C-trained continuous-weight networks. Red (or blue) numbers indicate the scores of EA that are higher than those of DQN (or A3C), but lower than A3C (or DQN).

| Game name | DQN,<br>continuous weight,<br>[69] | A3C,<br>continuous weight,<br>[69] | EA,<br>continuous weight,<br>[69] | EA,<br>binary weight,<br>ours | Hebbian-Q,<br>continuous weight,<br>ours |
| --- | --- | --- | --- | --- | --- |
| amidar | 978 | 264 | 263 | 251 | 10 |
| assault | 4,280 | 5,475 | 714 | 873 | 130 |
| asterix | 4,359 | 22,140 | 1,850 | 1,923 | 92 |
| asteroids | 1,365 | 4,475 | 1,661 | **1,596** | 146 |
| atlantis | 279,987 | 911,091 | 76,273 | 74,563 | 1,400 |
| enduro | 729 | -82 | 60 | **32** | 5 |
| frostbite | 797 | 191 | 4,536 | **4,520** | 30 |
| gravitar | 473 | 304 | 476 | **589** | 26 |
| kangaroo | 7,259 | 94 | 3,790 | **3,997** | 151 |
| seaquest | 5,861 | 2,355 | 798 | 800 | 85 |
| skiing | -13,062 | -10,911 | -6,502 | **-6,503** | -28,383 |
| venture | 163 | 23 | 969 | **1,070** | 22 |
| zaxxon | 5,363 | 24,622 | 6,180 | **6,195** | 24 |

## Discussion

Overall, by unifying a broad spectrum of biologically experimental evidence into a coherent picture (**Figures 1-5**), we show that heterosynaptic plasticity together with dendrite gating implies a biologically plausible EA mechanism of brain learning. Our EA model provides new insights into the origin of brain dynamics in cognitive tasks (**Figure 6**), and manifests its broad competence and powerful capability in training spiking or analog neural networks (**Figures 7, Table 1**). Compared to the homosynaptic mechanisms (i.e., local plasticity rules) reviewed in [3], our mechanism has the following computational advantages:

1. EA is model-free: it does not need information on the network architecture or task objective. Synaptic updating in our EA model is caused by spontaneous mutation; unlike in backpropagation algorithms, delicate feedback mechanisms are necessary to guide synaptic changes: this is the reason for the broad competence of our EA model for any network architecture, maximizing the generalizability of the model. Besides, a recent proposal of brain-learning mechanism suggests that evolution may endow each synapse with a specific set of parameters of Hebbian learning, so that the brain can learn in a manner that different synapses change in different Hebbian-type rules [70, 71]. In this proposal, the information on how each synapse should change for real-life tasks is coded in DNA. Though this proposal does not need backpropagated signals to guide synaptic updating, it requires infeasibly huge information of over 100 trillion synapses in the brain [72] coded in DNA. Our EA model, however, does not need much DNA-encoded knowledge.
2. EA is particularly good at creatively dealing with multi-objective optimization problems [73]. Real-world problems usually require addressing several conflicting objectives. For example, because a cheap house in downtown is usually unavailable, a house buyer may consider both an expensive house in the downtown and a cheap house in the suburb, due to their own merits to be either convenient or cheap. Gradient-based methods can solve this problem using a single objective, considering it as a linear combination of these multiple objectives with pre-assigned preferential weights: e.g., a house buyer may emphasize more on price or convenience before making the decision. In contrast, a population of agents trained by EA can simultaneously enumerate a number of possible solutions, each with its unique merit (more precisely, the Pareto front [73]), thereby significantly improving the creativity in dealing with conflicting objectives. In neuro-imaging studies, it has been found that the functional connectivity of the brain is dynamically reconfigured, and this reconfiguration is closely related to the creativity of human subjects [74, 75]. In terms of our model, this dynamic reconfiguration is caused by the alternation of activated agents (**Figure 1e**), reflecting the solution enumeration process mentioned above.
3. Natural selection creates our brain, endowing us with amazing learning capabilities, such as meta-learning [76], zero-shot (or few-shot) learning [77], transfer learning [78], continuous learning [79], etc. Therefore, EA is a universal philosophy to automatically create high intelligence in dealing with a complicated environment. By taking EA as its learning mechanism, the brain can potentially maximize its adaptation to the environment, facilitating the survival of the animal.

Compared with gradient-based methods, the drawback of EA is slow learning speed. Classification of the MNIST images requires over 10,000 learning epochs using EA (**Supplementary Figure 3b**), in contrast to dozens or hundreds of epochs using gradient-based methods. However, learning speed may not be a critical aspect, because EA mainly mediates the off-line learning during resting or sleeping (**Figure 4**), whose neuronal firing sequences are temporally compressed by about 20-fold relative to those in real experience [59]. Besides, the brain may use several strategies to speed up learning:

1. First is the combination of homosynaptic mechanisms that are compatible with gradient-based methods [3]. To manifest the sole power of heterosynaptic plasticity, we do not add any local homosynaptic rules in our model. Adding local homosynaptic rules can guide the mutation direction using hints of gradients, potentially speeding up learning.
2. Curriculum learning (i.e., learning the easier aspects of the task first and then gradually increasing the task difficulty) can significantly improve the speed and success rate of learning complicated tasks [80, 81]. To achieve optimal learning effects, the brain may adapt the contents of replay (so that the replayed contents are neither too simple nor too complex) to its current capability in solving problems: a process called autocurriculum [81]. It has been found that the brain prioritizes high-rewarded memories to replay [82], which may be the mechanism of autocurriculum: the replayed contents automatically adjust their complexity, and prioritize the highest-rewarded situations within the current problem-solving ability of the brain, training the brain to be more powerful.
3. A recent theoretical study [83] suggests that compared with EA with only mutation, EA with crossover operation is faster by polynomial order of the number of agents. Our model only contains the mutation operation, but can easily include crossover. Suppose the network *N* (**Figure 1d**) has two modules *N_A_* and *N_B_*, whose dendritic routes are separately controlled by two networks *A* and B, such that when a neuron *a_i_* ∈ *A* (or *b_j_* ∈ *B*) is activated, the dendritic routes labeled by *i* (or *j*) in *N_A_* (or *N_B_*) are gated on, with other routes gated off. If at the early stage of training, *a*_1_ and *b*_1_ (or *a*_2_ and *b*_2_) are activated simultaneously, such that dendritic routes with the same label 1 (or 2) in both *N_A_* and *N_B_* are gated on simultaneously; but at a later stage, *a*_1_ and *b*_2_ (or *a*_2_ and *b*_1_) are activated simultaneously, such that dendritic routes with different labels in *N_A_* and *N_B_* are gated on simultaneously: then this later stage fulfills crossover operation.

The most significant value of our work is to contribute a novel paradigm to the thought reservoir of neuroscience, so that future neuroscientists may think from the aspect of EA to interpret observations and make predictions, rather than only from the aspects of local rules and backpropagation. A testable prediction of our work is the importance of the dynamic reconfiguration of information routes in the neural-microcircuit level for learning (**Figure 1e**), particularly during off-line replay at rest or in sleep (**Figure 4**). In our model, this dynamic reconfiguration is realized by the alternation of dendritic gating by SST interneurons (**Figure 1e**), but it may also be realized by the alternative inhibitory shunting of different neuronal ensembles [84]. Our work also implies a potential optogenetic therapy for Alzheimer’s disease (AD). It has been found that the dynamic variability of the functional connectivity of AD human patients is reduced [85], and the neuronal firing in the hippocampus of AD mice follows rigid sequences, lacking the adaptation to tasks and environment in normal mice [86]. Therefore, AD is possibly related to the rigidness of information routes in the brain, and may be ameliorated if we optogenetically stimulate the SST interneurons using time-varying patterns. Optogenetic stimulation of PV interneurons is a promising therapy for AD [87]; its combination with the optogenetic stimulation of SST interneurons is an exciting research direction.

## Methods

### XOR task

In the network of **Figure 5c**, 20 pyramidal neurons receive from two groups of excitatory neurons (each group has 20 neurons) through 20 dendrites gated by different SST interneurons. These pyramidal neurons are inhibited by PV interneurons, which receive input from both groups of excitatory input neurons. Pyramidal neurons also give output to a single neuron. Pyramidal neurons, PV interneurons and the output neuron are modeled with leaky integrate-and-fire equations:

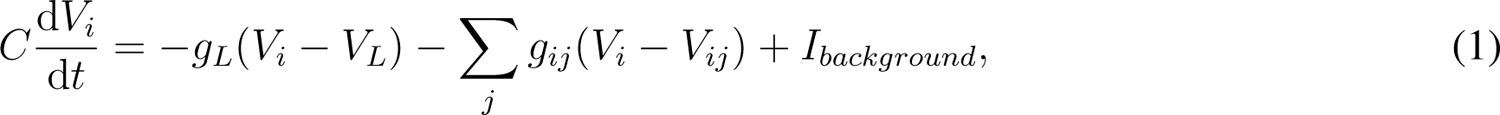

where *V_L_* = −74mV, *g_L_* = 25nS, *C* = 500pF, and *I_background_* = 425pA (parameters are chosen according to [10]). When the membrane potential *V_i_* reaches the firing threshold *V_θ_* = 54mV, a spike is recorded and *V_i_* is reset to *V_reset_* = −60mV. The reversal potential *V_ij_* is 0mV if the synapse from *j* to *i* is excitatory, or −70mV if this synapse is inhibitory. The dynamics of the synaptic conductance *g_ij_* is

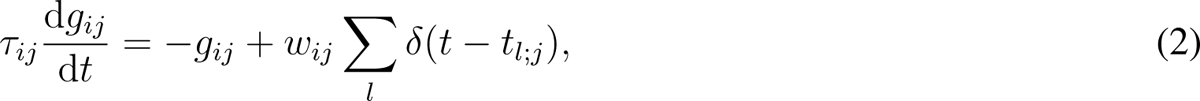

with *t_l_*_;_*_j_* being the time of the *l*th spike of the *j*th neuron. The time constant *τ_ij_* is 10ms for the synapses from the PV interneurons to the pyramidal neurons and 5ms for the other synapses. The synaptic efficacy *w_ij_* is 4.8nS from the excitatory input neurons to the PV interneurons, 2.4nS from the inhibitory input neurons to the PV interneurons, 2.4nS from the PV interneurons to the pyramidal neurons, and 4.8nS from the pyramidal neurons to the single output neuron. All the above synapses are connected with probability 0.4, and fixed during training. The synapses from the excitatory input neurons to the pyramidal neurons are all-to-all connected in every dendrite of the pyramidal neurons, and have either large efficacy of 2.4nS or small efficacy of 0nS. The excitatory input neurons produce Poisson spike trains. The firing rates of the two groups of excitatory input neurons are respectively (0Hz, 0Hz), (20Hz, 0Hz), (0Hz, 20Hz), and (20Hz, 20Hz) in the four successive intervals with duration 500ms in each training epoch (**Figure 5d, upper panel**). The pyramidal neurons also receive from a group of inhibitory neurons (not depicted in **Figure 5c**), which output Poisson spike trains of 40Hz. Our simulation was performed in the Brian simulator [88] using the exponential Euler method with time step of 0.1ms.

The potentiation eligibility trace of the synapse from a pre-synaptic excitatory input neuron to a post-synaptic pyramidal neuron through the *d*th dendrite is

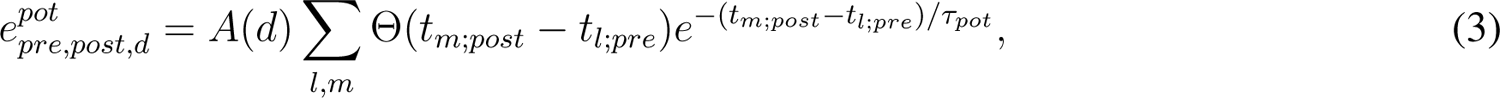

where Θ(·) is the step function, *t_m_*_;_*_post_* (or *t_l_*_;_*_pre_*) is the time of the *m*th (or *l*th) spike of the post- (or pre-) synaptic neuron, *τ_pos_* = 20ms is the characteristic time window indicating the simultaneity of the pre- and post-synaptic spikes that induces potentiation eligibility trace, *A*(*d*) = 1 for a synapse on the gated-on dendrite, and *A*(*d*) = 0.5 for a synapse on a gated-off dendrite modeling the effect of diffusive NO emitted from the gated-on dendrite. Here we suppose that the potentiation effect of the NO emitted from a gated-on dendrite of a pyramidal neuron can only influence the dendrites of the same pyramidal neuron, and has no influence on the dendrites of other pyramidal neurons, based on the experimental observation that the spatial scope of this potentiation effect is comparable with the dendritic arbor of a pyramidal neuron [15, 14].

Synaptic efficacy was updated according to the eligibility trace and the dopamine level, satisfying the following rules or constraints, explained below:

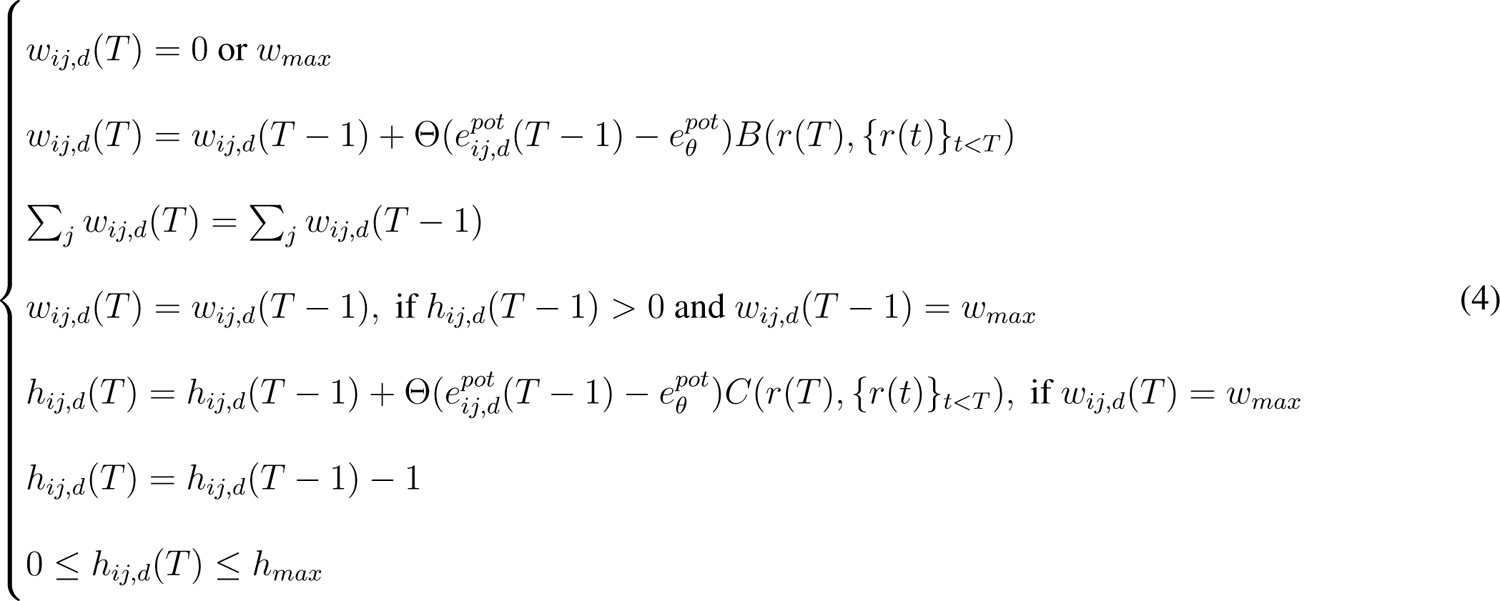

The first equation indicates that the synapse *w_ij,d_*(*T*) from the *j*th excitatory input neuron to the *i*th pyramidal neuron through the *d*th dendrite at the *T* th epoch has binary efficacies 0 or *w_max_* = 2.4nS. The second equation indicates that *w_ij,d_* is potentiated if *e^pot^_ij,d_* is larger than a threshold value *e^pot^_θ_* = 0.45 and the factor *B* controlled by the reward *r*(*T*) at the *T* th epoch is larger than zero. In our simulation, *B* = 1 if *r*(*T*) is not smaller than the second largest reward obtained in the last 20 training epochs, and *B* = 0 otherwise. The reward at an epoch is defined as

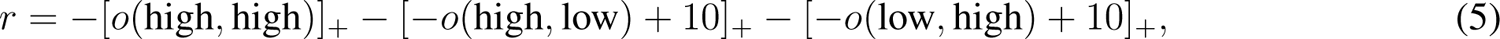

where *o*(high, low) means the number of spikes of the output neuron during the interval of 500ms when Group 1 of the excitatory input neurons have high firing rate and Group 2 have low firing rate; *o*(high, high) and *o*(low, high) have similar definitions; *o*(low, low) = 0 all the time. The third equation in Eqs. 4 indicates dendrite-level synaptic homeostasis (**Figure 3c, lower panel**), which keeps constant the total synaptic efficacy on a dendrite (specifically, half of the synapses on a dendrite are large). The fourth equation indicates that if the hidden state *h_ij,d_* in a synapse with large efficacy is larger than zero, this synapse will remain unchanged in the current epoch. In other words, a large synapse can only be depressed when its hidden state is zero. Note that the hidden state remains at zero for synapses with low efficacy. The fifth equation indicates that if the eligibility trace *e^pot^* is larger than *e^pot^* and the factor *C* controlled by the reward is larger than zero, the hidden state in the large synapse will be increased. In our simulation, *C* is 20 if *r*(*T*) is not smaller than the largest reward obtained in the last 20 training epochs, *C* is 10 if *r*(*T*) is smaller than the largest but not smaller than the second largest reward obtained in the last 20 training epochs, and *C* = 0 otherwise. The sixth equation indicates that the hidden state in every large synapse is decreased by 1 in each training epoch, so that the synapse gradually becomes less stable. The seventh equation indicates that the hidden state is non-negative and no larger than a maximum value *h_max_* = 20.

The second and third equations in Eqs. 4 may conflict with each other. In our simulation, when too many synapses had large potentiation eligibility traces and were to be potentiated, we kept synaptic homeostasis (i.e., the third equation) non-violated, and set the synapses with largest potentiation eligibility traces at large efficacy. Specifically, at the *T* th epoch when *B >* 0, we collected the synapses either with *w_ij,d_*(*T* − 1) *>* 0 and *h_ij,d_*(*T* − 1) = 0 or with *w_ij,d_*(*T* − 1) = 0 and *e^pot^_ij,d_* (*T* − 1) *> e^pot^_θ_* into a set *S*, sorted the potentiation eligibility traces in the synapses in S in descending order, and let the first *N*_input_*/*2 − *N*_stable large_ synapses have large efficacy, and let the rest synapses have small efficacy. Here, *N*_input_ = 40 is the number of synapses on a single dendrite (which is the total number of excitatory neurons in the two input groups), and we constrained the number of synapses with large efficacy on a dendrite fixed at *N*_input_*/*2, modeling dendrite-level synaptic homeostasis (the third equation of Eqs. 4). *N*_stable large_ is the number of synapses with *w_ij,d_*(*T* − 1) *>* 0 and *h_ij,d_*(*T* − 1) *>* 0 on the *d*th dendrite.

In our model, each pyramidal neuron has 20 dendrites, labeled from 1 to 20. Every neuron in the two input populations gives output to every dendrite (**Figure 5c**). At each training epoch, the dendrites of all the pyramidal neurons with the same randomly selected label were gated-on, with the other dendrites gated off (**Figure 1e**). Synaptic mutation was performed at the gated-on dendrites at the beginning of each epoch. In individual mutation, synapses with large efficacy and zero hidden state were randomly set to small efficacy with probability 0.25 at the gated-on dendrites; at the same time, the same number of randomly selected synapses with small efficacy were also set to large efficacy, to maintain the total number of large synapses on every gated-on dendrite. In group mutation, all the synapses (except for those synapses with non-zero hidden states) from a randomly selected input group to the gated-on dendrite of a randomly selected pyramidal neuron were set to small efficacy; at the same time, the same number of synapses on the gated-on dendrite from another input group to the selected pyramidal neuron were set to large efficacy to maintain the total number of large synapses on that gated-on dendrite. In both mutation strategies, a single large synapse with zero hidden state has almost equal probability 0.025 to be mutated (**Supplementary Figure 1c**), so that the performance difference under these two strategies (**Figure 5e, Supplementary Figure 1b, c**) is caused by these strategies themselves, instead of the difference of mutation rate.

### Context-dependent decision-making task

Our model to study the context-dependent decision-making task in **Figure 6** closely follows that of [28]. The recurrent neural network contains *N* = 100 neurons, and is all-to-all connected with the following dynamics:

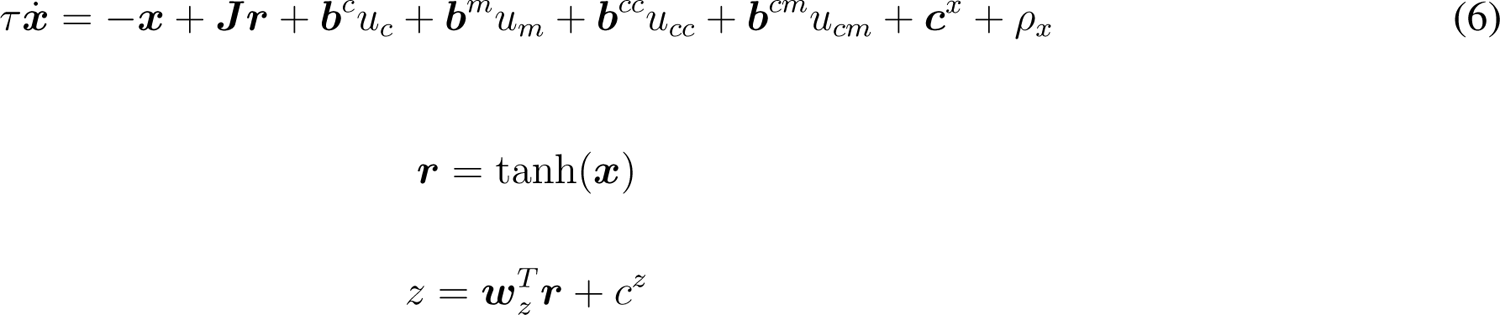

The variable ***x***(*t*) is a *N*-dimensional vector containing the total synaptic current of each neuron in the network, and ***r***(*t*) are the corresponding firing rates. Each neuron has a time constant *τ* = 10ms. The matrix ***J*** defines the recurrent connections in the network. The network receives 4-dimensional input, ***u***(*t*) = [*u_c_*(*t*)*, u_m_*(*t*)*, u_cc_*(*t*)*, u_cm_*(*t*)]*^T^*, through synaptic weights, ***B*** = [***b****^c^,* ***b****^m^,* ***b****^cc^,* ***b****^cm^*]. These four inputs represent, respectively, the sensory evidence for color and motion, and the contextual cues instructing the network to integrate either the color or the motion input. Finally, ***c****^x^* is a vector of offset currents and *ρ_x_* is a white noise drawn at each time step with standard deviation 0.1. The output *z* of the network is a weighted sum of the firing rates, with weights ***w****^T^* and bias *c^z^*. All the synaptic connections in ***J***, ***B*** and ***w****_z_* are binary, taking values of ±1*/*√*N*, ±1 and ±1*/*√*N*, respectively. During training, the network dynamics were integrated for the duration *T* = 750 ms using Euler updates with Δ*t* = 1ms. After training, model dynamics were integrated for an additional 200 ms with the sensory inputs turned off, according to which we plotted the orange dots and lines in **Figure 6c, d**.

The contextual inputs *u_cm_* and *u_cc_* were constant for the duration of the trial. In the motion context *u_cm_*(*t*) = 1 and *u_cc_*(*t*) = 0, while in the color context *u_cm_*(*t*) = 0 and *u_cc_*(*t*) = 1. The motion and color inputs *u_m_* and *u_c_* are one-dimensional white-noise signals:

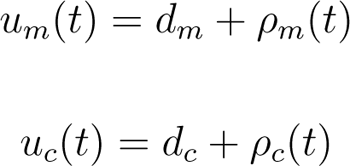

The white noise terms *ρ_m_* and *ρ_c_* have zero mean and standard deviation 1. During training, the offsets *d_m_* and *d_c_* were randomly chosen on each trial from the range [-0.1875, 0.1875]. During simulations after training, *d_m_* and *d_c_* took 6 values (±0.009, ±0.036, ±0.15), corresponding to weak, intermediate, and strong evidence toward either choice. In the psychometric curves (**Figure 6b**), the coherence value is normalized so that ±0.15 of *d_m_* or *d_c_* corresponds to ±1 of the horizontal coordinate in the figure.

The target *p* of network training is that the output *z* approaches 1 (or −1) at time *T* when *d_m_ >* 0 (or *<* 0) in the motion context, and approaches 1 (or −1) when *d_c_ >* 0 (or *<* 0) in the color context. In the EA, the number of agents was 100, the number of elite agents in each generation was 5. These agents were evaluated by averaging (*p* − *z*(*T*))^2^ over 128 simulation trials. At each training epoch, every non-parental agent copied the synaptic weights ***J***, ***B*** and ***w****_z_* as well as the bias ***c****^x^* and *c^z^* from a randomly chosen elite agent, with the following mutation: every synaptic weight was flipped with probability 0.0005; to every bias was added a random Gaussian number with zero mean and standard deviation 0.002. The mutation of bias models the mutation of synaptic weights inputted from other brain areas.

The axes of choice, motion and color in **Figure 6c, d** were obtained by first linear regressing the trajectory ***x***(*t*) of all simulation trials and then orthogonalizing the regression coefficient using QR-decomposition [28]. Similar to [28], we plotted ***x***(*t*) − ⟨***x***(*t*)⟩_trial_ in **Figure 6c, d**, with the urgency signal ⟨***x***(*t*)⟩_trial_ being the average trajectory over all simulation trials in the motion (**Figure 6c**) or color (**Figure 6d**) context.

The attractors in **Figure 6c-e** were obtained by minimizing the function

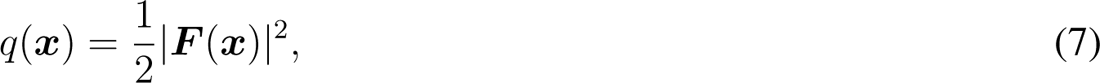

where ***F*** (***x***) is the right-hand side of eq. 6 after setting *u_c_* = *u_m_* = *ρ_x_* = 0, representing the speed of changing of neural state. To get the attractors in the motion context, *u_cm_*(*t*) = 1 and *u_cc_*(*t*) = 0; to get the attractors in the color context, *u_cm_*(*t*) = 0 and *u_cc_*(*t*) = 1. We used the L-BFGS-B algorithm of the ‘minimize’ routine of the Scipy package to minimize *q*(***x***), initializing ***x*** to be a random point in the trajectory of ***x***(*t*). To find slow-dynamic points on the line attractor instead of the two stable fixed points, we set the tolerance parameter ‘ftol=0.002’ to early stop the minimization algorithm. The found slow-dynamic points have a close-to-zero negative eigenvalue and many large negative eigenvalues. The small bars in **Figure 6e** are the left eigenvectors of the close-to-zero eigenvalues of these points (i.e., selection vectors), projected in the subspace spanned by the input weights ***b****^c^* and ***b****^m^*.

### Trajectory generation task

In the trajectory generation task (**Figure 7a, b**), we considered a network of *N* = 1000 randomly and sparsely connected (connection probability *p* = 0.1) quadratic integrate-and-fire neurons with dynamics [89]

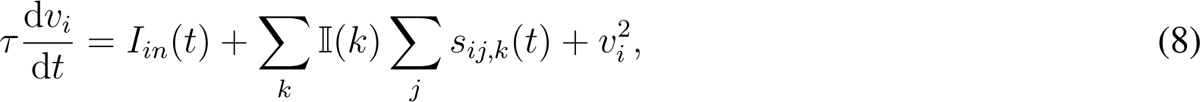

where *τ* = 10ms; *v_i_* is a dimensionless variable representing the membrane potential; *s_ij,k_* is the synaptic current from neuron *j* to neuron *i* through dendrite *k*; and I(*k*) = 1 or 0, indicating whether or not dendrite *k* is gated on. The dynamics of *s_ij,k_* is

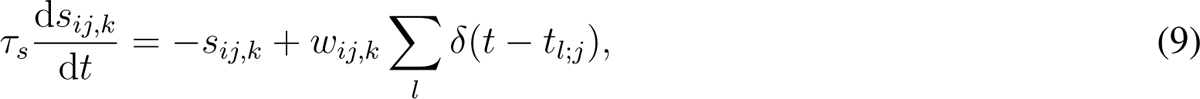

where *τ_s_* = 20ms, *w_ij,k_* is the synaptic weight from neuron *j* to neuron *i* through dendrite *k*, and *t_l_*_;_*_j_* is the time of the *l*th spike of neuron *j*. To simulate the dynamics of quadratic integrate-and-fire neurons, we used theta neuron model derived by a simple change of variables *v_i_* = tan(*θ_i_/*2), getting

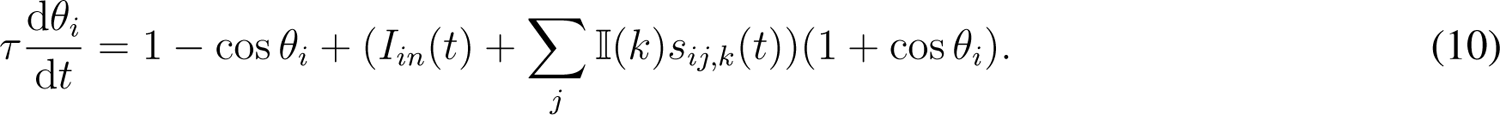

The output of the network is

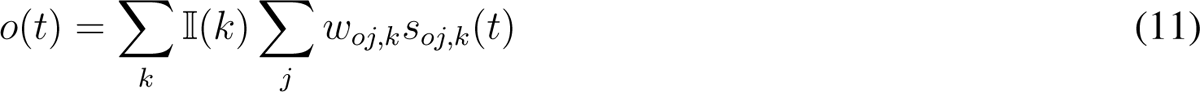

where *w_oj,k_*is the output weight of neuron *j* througth dendrite *k*, *s_oj,k_* is the synaptic current with the dynamics

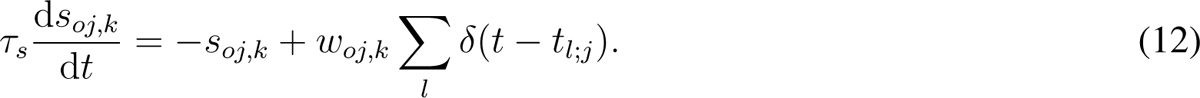

The recurrent synapse *w_ij,k_* takes binary values: either 20*/*√*pN* or −20*/*√*pN*. The output synapse *w_oj,k_* also takes binary values: either either 1*/*√*N* or −1*/*√*N*.

The target trajectory (orange curve in **Figure 7b**) was defined as *f* (*t*) = *A* sin(2*π*(*t* − *T*_0_)*/T*_1_) sin(2*π*(*t* − *T*_0_)*/T*_2_), where *A*, *T*_0_, *T*_1_ and *T*_2_ were randomly sampled from intervals [0.5, 1.5], [0, 1000 ms],[500 ms, 1000 ms] and [100 ms, 500 ms respectively. At the beginning of each simulation session, every neuron was stimulated for 50 ms with constant external stimulus that had random amplitude sampled from [−1, 1] (red shading in **Figure 7b**). The pattern of the stimulation was the same for the same target function. Simulations were performed in the Brian simulator [88] using the Euler method with time step of 0.1ms.

Our EA was performed with the aim to reduce the loss function

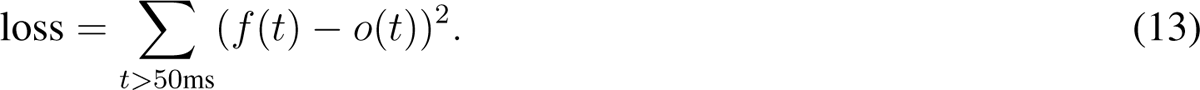

In the EA, the number of agent was 100, the number of elite agents in each generation was 5. At each training step, every non-parental agent copied the synaptic weights of a randomly chosen elite agent, then every synaptic weight was flipped with probability 0.0005.

### MNIST classification task

In **Figure 7c, d**, we train a multi-layer perceptron with 2 hidden layers with 256 and 128 neurons respectively to classify the MNIST images (size 28 × 28 = 784 pixels) into 10 classes. Adjacent layers in the feedforward network are all-to-all connected. The non-linear activation function is ReLU.

The EA was performed on networks whose synaptic weights took binary values {−1*/*√*N*_in_,1*/*√*N*_in_}, where *N*_in_ represents input size: 784 for weights between the input layer and the first hidden layer; 256 for weights between the first and second hidden layers; and 128 between the second hidden layer and the output layer. We initialized 100 network configurations. At every training epoch, negative log-likelihood loss averaged over all the 60000 images in the training dataset was evaluated for every configuration. The elite configuration (the one with the smallest loss) remained unchanged, whereas each weight of the other configurations was updated to be the corresponding weight of the elite configuration with probability 0.99995, and to be different with probability 0.00005 (mutation). Therefore, there is only one parent in each generation of agents in this algorithm. We also studied the case of dendrite-level synaptic homeostasis (**Figure 3c, lower panel**), in which the number of synapses with large efficacy received by a neuron in a configuration was kept fixed after the random initialization of synaptic weights. Adding this homeostasis mechanism hardly influences the classification performance (**Supplementary Figure 3b**).

Back-propagation was performed on networks whose synaptic weights took continuous values. We used Adam optimizer of Pytorch for the training. Batch size was 100, learning rate was 0.0001, the number of training epochs was 400.

The Hebbian learning was also performed on continuous-weight networks. All the low-level layers (except the top layer) were trained by competitive Hebbian learning [65]. In every step, the weights were updated by

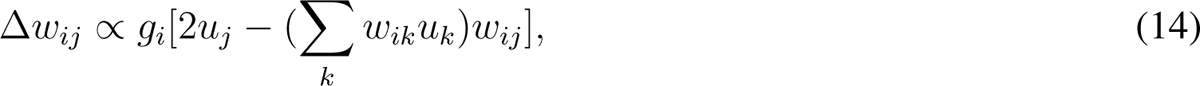

where *w_ij_* is the synaptic weights from the *j*th pre-synaptic neuron to the *i*th post-synaptic neuron, *u_j_* is the input from the *j*th pre-synaptic neuron, *g_i_* is 1 for the highest activated post-synaptic neuron, −0.4 for the second highest activated post-synaptic neuron, and 0 for others. Batch normalization was performed on the pre-synaptic input *u_j_*. Layers were trained one after another, such that when a higher-level layer was trained, all the lower-level layers were fixed. Adam optimizer was used to supervisedly train the top layer. Following [65], the learning rate of Hebbian learning linearly decreased from the maximal value 0.04 at the first epoch to 0 at the last epoch, and the learning rate of supervised learning was kept at 0.0001. Both Hebbian and supervised training was performed with minibatch size 100 for 1000 epochs.

### Atari game task

The structure of the deep neural network to play Atari games closely followed those of [67] and [69]. Specifically, the neural network mapped 4 recent frames of size 84 × 84 to actions through 3 convolutional layers and 2 fully-connected layers. Of the three convolutional layers, the kernel sizes were 8 × 8, 4 × 4 and 3 × 3 respectively, the strides were 4, 2 and 1 respectively, and the numbers of features were 32, 64 and 64 respectively. The first fully-connected layers had 512 neurons. ReLU activation function was used.

EA was used to train neural networks whose synaptic weights took binary values {−2*/*√*N*_in_,2*/*√*N*_in_}, with *N*_in_ being the input size. The training protocol closely followed that of [69]. Specifically, each generation had 1000 agents, each was evaluated by one episode (i.e., from the start to the end of a game, when the player succeeded or was killed in the game). The top 20 agents were selected to be parents and reproduced the next generation by flipping their binary weights with probability 0.002 (mutations). Then the top 10 agents were further evaluated by 30 episodes, based on which the elite agent was selected to be the agent that achieved the highest score. The elite agent became a member of the next generation without any mutation. The performance of EA at any training epoch was the average score of the elite agent in further 200 episodes. We trained the network in 2.5 × 10^8^ epochs. Each number of EA in **Table 1** represents the median value of the scores in the final epochs of 5 training trials.

To perform Hebbian-Q learning, batch normalization was inserted between each layer of the network. The low-level layers were trained using Eq. 14, and the last layer was trained supervisedly using Adam optimizer. When training a higher-level layer, all lower-level layers were kept fixed.

When unsupervisedly training a low-level layer, at each time step, a random action was taken of a game [90], then minibatches of size 32 were randomly extracted from a buffer of 10^6^ recent frames to train the network using Eq. 14. To train convolutional layers, we averaged over the weight updating at different spatial locations to update the sharing weights. Similar to [65], learning rate linearly decreased from the maximal value 0.02 at the first epoch to 0 at the last epoch. The three convolutional layers were trained in 1.6 × 10^5^ epochs, and the first fully-connected layer was trained in 4.8 × 10^5^ epochs.

When supervisedly training the top layer, minibatches of size 32 were randomly extracted from a buffer of 10^6^ recent frames at each epoch in a similar protocol to [67]. The action policy during training was *ϵ*-greedy, with *ϵ* linearly annealed from 1 to 0.1 in the first 10^6^ epochs. The target network was cloned from the Q-network every 10000 epochs. Every 2 × 10^5^ epochs, the performance of the agent was evaluated by the average score over 200 episodes of games. We trained the top layer for 2 × 10^6^ epochs, and found no sign of continuing increase of performance. Each number of Hebbian-Q in **Table 1** represents the maximal value of the evaluation scores in 2 training trials.

## Acknowledgement

This work was supported by the National Natural Science Foundation of China (Grant no. 32000694 to Z. B., Grant no. 12175242 to D. Y. and Grant no. 32071141 to Y. Z.), the Natural Science Foundation of Shandong province (Grant no. ZR2019ZD34 to Y. Z.) and the start-up fund of the Institute for Future of Qingdao University (to Z. B.).

---

**Supplementary Figure 1:**
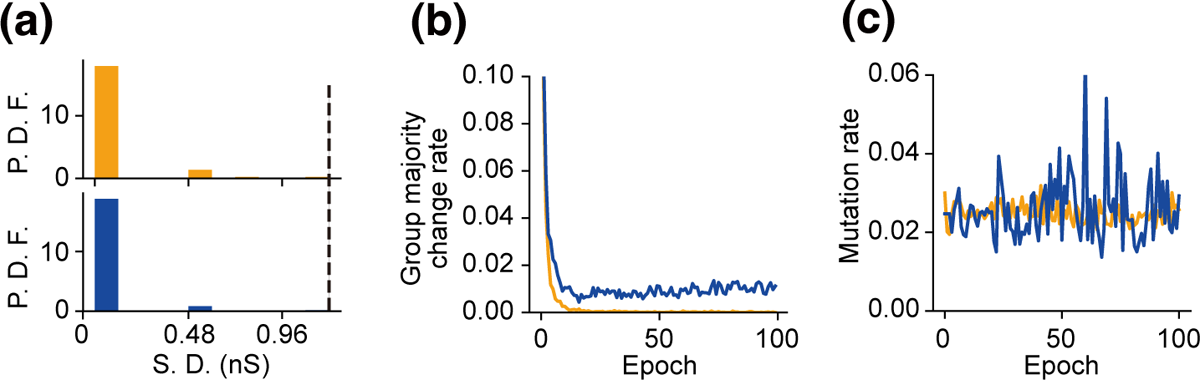
Cooperative plasticity of the synaptic weights in the XOR task. (**a**) Probability density function (P. D. F.) of the standard deviation (S. D.) of {*w_d,i_*}*_i∈Gk_*, with {*w_d,i_*}*_i∈Gk_* (*k* = 1, 2) being the efficacies of all the synapses the neurons in group *G_k_* through a given dendrite *d*, under individual mutation (orange) and group mutation (blue). The dashed vertical line indicates the S. D. when the synapses in {*w_d,i_*}*_i∈Gk_* have either large or small efficacies with equal probability. This panel means that the synapses from the same group either mostly have large efficacy or mostly have small efficacy. (**b**) The probability that the majority value of {*w_d,i_*}*_i∈Gk_* changes at a given epoch during the training process. This panel means that the majority value of {*w_d,i_*}*_i∈Gk_* frequently changes under group mutation, but hardly changes under individual mutation, consistent with Figure 5a**, b**. (**c**) The mutation rate of large synapses with zero hidden states under individual mutation (orange) or group mutation (blue) as a function of training epoch. Notice that they are almost the same at our parameter values, so the performance difference under these two mutation strategies (Figure 5e) is due to these strategies themselves rather than the difference of mutation rate. The results in the three panels are average over 100 training trials.

**Supplementary Figure 2:**
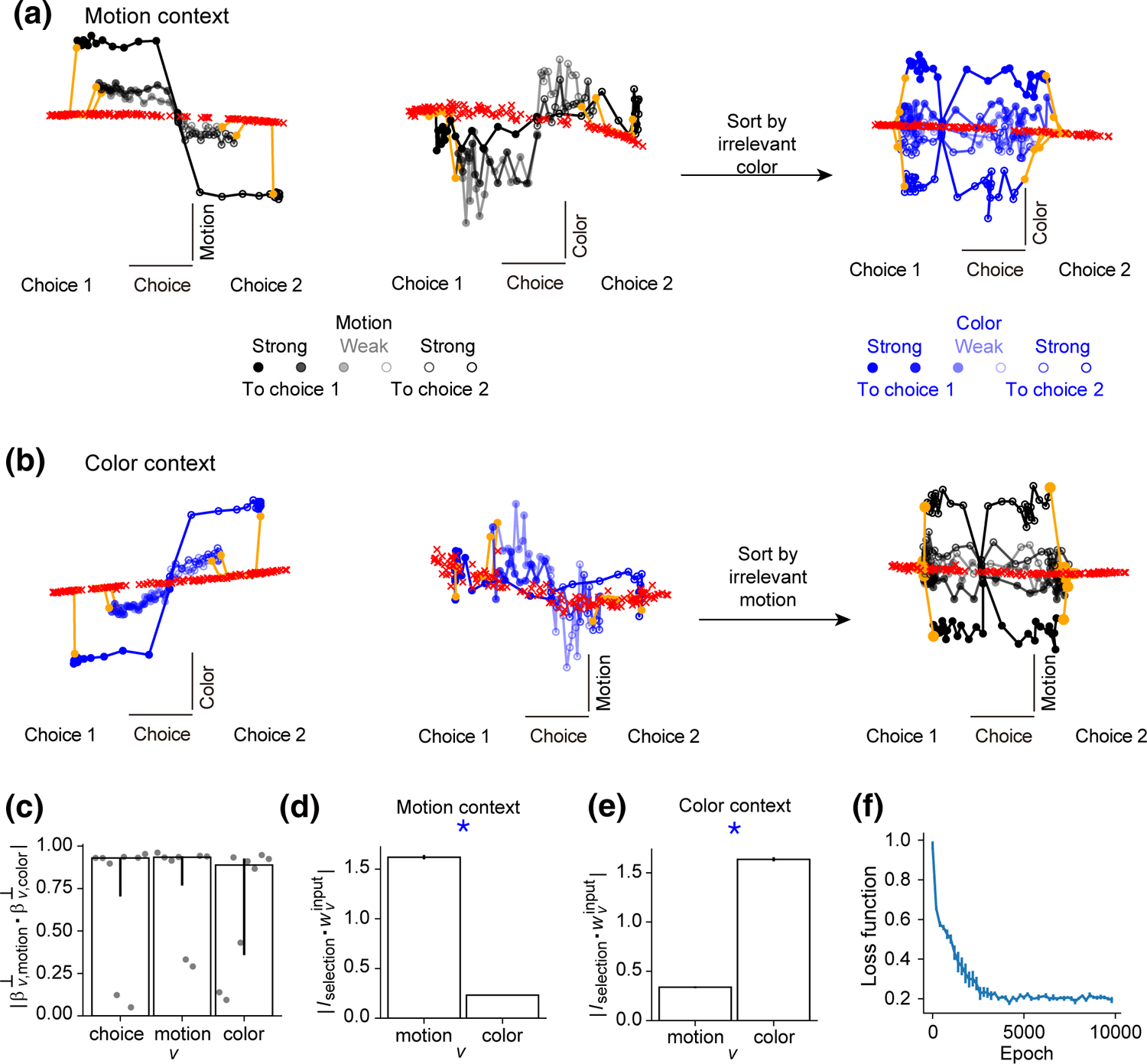
The dynamics of EA-trained neural networks in the context-dependent decision-making task. (**a**) Left: Dynamics of model population responses in the motion context in the subspace spanned by the axes of choice and motion, when the motion coherence takes different values. Middle: Same data as the left panel, but in the subspace spanned by the axes of choice and color. Right: Same data as the middle panel, but re-sorted according to the direction and strength of the irrelevant color input. (**b**) Responses in the color context, analogous to panel **a**. (**c**) *β^⊥^_v, motion_* (*v* =choice, motion, color) indicates the unit vector along the choice, motion or color axis in the motion context; *β^⊥^_v, color_* indicates the unit vector in the color context. This panel means that the choice, motion or color axes in different contexts are almost colinear. Error bars represent quantiles over 8 simulation trials. (**d**) Motion context. *w*^input^_v_ (*v* =motion, color) is the input weight of the motion or color signal; *I*_selection_ is the selection vector of the fixed points along the line attractor. This panel means that the motion input has a larger projection on the selection vector than the color input in the motion context. Error bars represent s.e.m. over 1024 fixed points found in 8 simulation trials. (**e**) Similar to panel **d**, but in the color context. This panel means that the color input has a larger projection on the selection vector than the motion input in the color context. (**f**) The loss function during the training process. Error bars represent s.e.m. over 8 simulation trials.

**Supplementary Figure 3:**
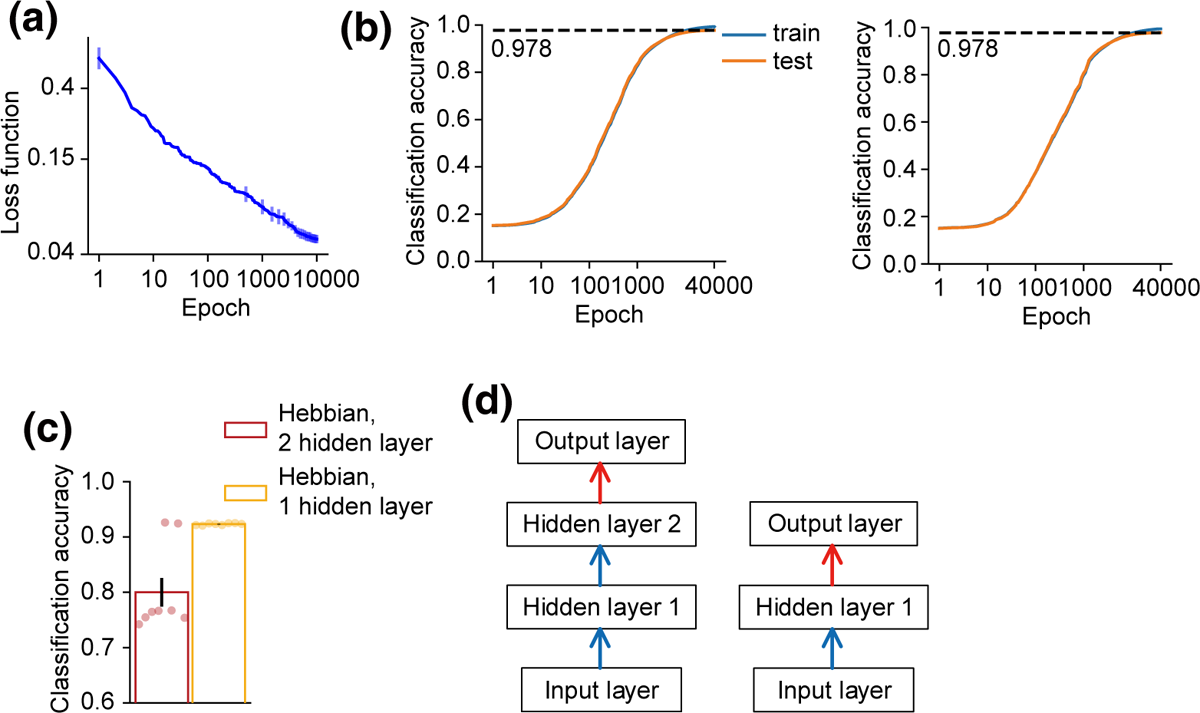
The trajectory generation task and the MNIST classification task. (**a**) Trajectory generation task. The loss function during the training process. (**b-d**) MNIST classification task. (**b**) Left: Classification accuracy during the training process of EA. The horizontal dashed line with the number indicates the final accuracy on the test dataset. Right: The same as the left panel, except under dendrite-level homeostasis. Notice that dendrite-level homeostasis hardly influences performance. (**c**) Classification accuracy on the test dataset when training deep networks with 2 hidden layers or 1 hidden layer using Hebbian rule. (**d**) Schematic of the structures of the networks examined in panel **c**, one with two hidden layers and the other with one hidden layer. Lower layers were trained using the Hebbian algorithm (blue arrows), and the top layer was trained by gradient-descent algorithm supervisedly (red arrows). In panels **a** and **c**, error bars represent s.e.m. over 8 training trials. Error bars in panel **b** (not shown) are comparable with the width of the lines.

## References

[1] Richards, B. A. et al. A deep learning framework for neuroscience. Nat. Neurosci. 22, 1761–1770 (2019).

[2] Dayan, P. & Abbott, L. F. Theoretical neuroscience: computational and mathematical modeling of neural systems (The MIT Press, Cambridge, 2001).

[3] Lillicrap, T. P., Santoro, A., Marris, L., Akerman, C. J. & Hinton, G. Backpropagation and the brain. Nat. Rev. Neurosci. 21, 335–346 (2020).

[4] Lillicrap, T. P., Cownden, D., Tweed, D. B. & Akerman, C. J. Random synaptic feedback weights support error backpropagation for deep learning. Nat. Commun. 7, 13276 (2016).

[5] Payeur, A., Guerguiev, J., Zenke, F., Richards, B. A. & Naud, R. Burst-dependent synaptic plasticity can coordinate learning in hierarchical circuits. Nat. Neurosci. 24, 1010–1019 (2021).

[6] LeCun, Y. Learning processes in an asymmetric threshold network. In Fogelman-Soulié, F., Bienenstock, E. & Weisbuch, G. (eds.) Disordered Systems and Biological Organization, 233–240 (Springer-Verlag, Les Houches, 1986).

[7] Lee, D.-H., Zhang, S., Fischer, A. & Bengio, Y. Difference target propagation. In Joint Eur. Conf. Machine Learning Knowl. Discov. Databases (2015).

[8] Fiete, I. R. & Seung, H. S. Gradient learning in spiking neural networks by dynamic perturbation of conductances. Phys. Rev. Lett. 97, 048104 (2006).

[9] Miconi, T. Biologically plausible learning in recurrent neural networks reproduces neural dynamics observed during cognitive tasks. eLife 97, e20899 (2017).

[10] Seung, H. S. Learning in spiking neural networks by reinforcement of stochastic synaptic transmission. Neuron 40, 1063–1073 (2003).

[11] Stanley, K. O., Clune, J., Lehman, J. & Miikkulainen, R. Designing neural networks through neuroevolution. *Nat*. Mach. Intell. 1, 24–35 (2019).

[12] Chistiakova, M., Bannon, N. M., Bazhenov, M. & Volgushev, M. Heterosynaptic plasticity: Multiple mechanisms and multiple roles. Neuroscientist 263, 532–536 (2014).

[13] Chistiakova, M. & Volgushev, M. Heterosynaptic plasticity in the neocortex. Exp Brain Res. 199, 377–390 (2009).

[14] Schuman, E. M. & Madison, D. V. Locally distributed synaptic potentiation in the hippocampus. Science 263, 532–536 (1994).

[15] Bonhoeffer, T., Staiger, V. & Aertsen, A. Synaptic plasticity in rat hippocampal slice cultures: local “Hebbian” conjunction of pre- and postsynaptic stimulation leads to distributed synaptic enhancement. Proc. Nat Acad. Sci. USA 86, 8113–8117 (1989).

[16] Chen, J. et al. Heterosynaptic long-term depression mediated by ATP released from astrocytes. Glia 61, 178–191 (2013).

[17] Deitmer, J. W., Verkhratsky, A. J. & Lohr, C. Calcium signalling in glial cells. Cell Calcium 24, 405–416 (1998).

[18] Chasse, R., Malyshev, A., Fitch, R. H. & Volgushev, M. Altered heterosynaptic plasticity impairs visual discrimination learning in Adenosine A1 receptor knock-out mice. J. Neurosci. 41, 4631–4640 (2021).

[19] Kepecs, A. & Fishell, G. Interneuron cell types are fit to function. Nature 505, 318–326 (2014).

[20] O’Connor, D. H., Wittenberg, G. M. & Wang, S. S.-H. Graded bidirectional synaptic plasticity is composed of switch-like unitary events. Proc. Natl Acad. Sci. USA 102, 9679–9684 (2005).

[21] Roelfsema, P. R. & Holtmaat, A. Control of synaptic plasticity in deep cortical networks. Nat. Rev. Neurosci. 19, 166–180 (2018).

[22] Duszkiewicz, A. J., McNamara, C. G., Takeuchi, T. & Genzel, L. Novelty and dopaminergic modulation of memory persistence: A tale of two systems. Trends Neurosci. 42, 102–114 (2019).

[23] Hulme, S. R., Jones, O. D., Raymond, C. R., Sah, P. & Abraham, W. C. Mechanisms of heterosynaptic metaplasticity. Phil. Trans. R. Soc. B 369, 20130148 (2013).

[24] Ji, D. & Wilson, M. Coordinated memory replay in the visual cortex and hippocampus during sleep. Nat. Neurosci. 10, 100–107 (2007).

[25] Harvey, C. D. & Svoboda, K. Locally dynamic synaptic learning rules in pyramidal neuron dendrites. Nature 450, 1195–1200 (2007).

[26] Hayama, T. et al. GABA promotes the competitive selection of dendritic spines by controlling local Ca2+ signaling. Nat. Neurosci. 16, 1409–1416 (2013).

[27] Engert, F. & Bonhoeffer, T. Synapse specificity of long-term potentiation breaks down at short distances. Nature 388, 279–284 (1997).

[28] Mante, V., Sussillo, D., Shenoy, K. V. & Newsome, W. T. Context-dependent computation by recurrent dynamics in prefrontal cortex. Nature 503, 78–84 (2013).

[29] Hiratani, N. & Fukai, T. Redundancy in synaptic connections enables neurons to learn optimally. Proc. Nat Acad. Sci. USA 115, E6871–E6879 (2018).

[30] Oldham, S. & Fornito, A. The development of brain network hubs. Dev. Cogn. Neurosci. 36, 100607 (2019).

[31] Wall, N. R. et al. Brain-wide maps of synaptic input to cortical interneurons. J. Neurosci. 36, 4000–4009 (2016).

[32] Pfeffer, C. K., Xue, M., He, M., Huang, Z. J. & Scanziani, M. Inhibition of inhibition in visual cortex: the logic of connections between molecularly distinct interneurons. Nat. Neurosci. 16, 1068–1076 (2013).

[33] Stott, J. J. & Redish, A. D. Representations of value in the brain: An embarrassment of riches? PLoS Biol. 13, e1002174 (2015).

[34] Fischer, A. G. & Ullsperger, M. An update on the role of serotonin and its interplay with dopamine for reward. Front. Hum. Neurosci. 11, e1004638 (2017).

[35] Mongillo, G., Rumpel, S. & Loewenstein, Y. Intrinsic volatility of synaptic connections - a challenge to the synaptic trace theory of memory. Curr. Opin. Neurobiol. 46, 7–13 (2017).

[36] Faisal, A. A., Selen, L. P. J. & Wolpert, D. M. Noise in the nervous system. Nat. Rev. Neurosci. 9, 292–303 (2008).

[37] Gerstner, W., Lehmann, M., Liakoni, V., Corneil, D. & Brea, J. Eligibility traces and plasticity on behavioral time scales: Experimental support of neohebbian three-factor learning rules. Front. Neural Circuits 12, 53 (2018).

[38] Schuman, E. M. & Madison, D. V. A requirement for the intercellular messenger nitric oxide in long-term potentiation. Science 254, 1503–1506 (1991).

[39] Kossel, A., Bonhoeffer, T. & Bolz, J. Non-hebbian synapses in rat visual cortex. Neuroreport 1, 115–118 (1990).

[40] Cornell-Bell, A. H., Finkbeiner, S. M., Cooper, M. S. & Smith, S. J. Glutamate induces calcium waves in cultured astrocytes: Long-range glial signaling. Science 247, 470–473 (1990).

[41] Scanziani, M., Malenka, R. C. & Nicoll, R. A. Role of intercellular interactions in heterosynaptic long-term depression. Nature 380, 446–450 (1996).

[42] Min, R. & Nevian, T. Astrocyte signaling controls spike timing-dependent depression at neocortical synapses. Nat. Neurosci. 15, 746–753 (2012).

[43] Andersson, M., Blomstrand, F. & Hanse, E. Astrocytes play a critical role in transient heterosynaptic depression in the rat hippocampal CA1 region. J. Physiol. 585, 843–852 (2007).

[44] Lee, C. M., Stoelzel, C., Chistiakova, M. & Volgushev, M. Heterosynaptic plasticity induced by intracellular tetanization in layer 2/3 pyramidal neurons in rat auditory cortex. J. Physiol. 590, 2253–2271 (2012).

[45] Tsumoto, T. & Suda, K. Cross-depression: an electrophysiological manifestation of binocular competition in the developing visual cortex. Brain Res. 168, 190–194 (1979).

[46] Takahashi, S. & Sakurai, Y. Sub-millisecond firing synchrony of closely neighboring pyramidal neurons in hippocampal ca1 of rats during delayed non-matching to sample task. Front. Neural Circuits 3, 9 (2009).

[47] Mirjalili, S. *Evolutionary Algorithms and Neural Networks: Theory and Applications* (Springer, Switzerland, 2019).

[48] Gao, S. et al. Dendritic neuron model with effective learning algorithms for classification, approximation, and prediction. IEEE Trans. Neural Netw. Learn. Syst. 30, 601–614 (2019).

[49] Holtmaat, A. & Caroni, P. Functional and structural underpinnings of neuronal assembly formation in learning. Nat. Neurosci. 19, 1–10 (2016).

[50] Josselyn, S. A. & Tonegawa, S. Memory engrams: Recalling the past and imagining the future. Science 367, eaaw4325 (2020).

[51] Abraham, W. C., Mason-Parker, S. E., Bear, M. F., Webb, S. & Tate, W. P. Heterosynaptic metaplasticity in the hippocampus in vivo: a BCM-like modifiable threshold for LTP. Proc. Natl. Acad. Sci. USA 98, 10924–10929 (2001).

[52] Hulme, S. R., Jones, O. D., Ireland, D. R. & Abraham, W. C. Calcium-dependent but action potential independent BCM-like metaplasticity in the hippocampus. J. Neurosci. 32, 6785–6794 (2012).

[53] Sutton, M. A. et al. Miniature neurotransmission stabilizes synaptic function via tonic suppression of local dendritic protein synthesis. Cell 125, 785–799 (2006).

[54] Barnes, S. J. et al. Deprivation-induced homeostatic spine scaling in vivo is localized to dendritic branches that have undergone recent spine loss. Neuron 96, 871–882 (2017).

[55] Royer, S. & Paré, D. Conservation of total synaptic weight through balanced synaptic depression and potentiation. Nature 422, 518–522 (2003).

[56] White, G., Levy, W. B. & Steward, O. Spatial overlap between populations of synapses determines the extent of their associative interaction during the induction of long-term potentiation and depression. J. Neurophysiol. 64, 1186–1198 (1990).

[57] Turrigiano, G. Too many cooks? Intrinsic and synaptic homeostatic mechanisms in cortical circuit refinement. Annu. Rev. Neurosci. 34, 89–103 (2011).

[58] Wolpert, D. & Macready, W. No free lunch theorems for optimization. IEEE Trans. Evol. Comput. 1, 67–82 (1997).

[59] Buch, E. R., Claudino, L., Quentin, R., Bönstrup, M. & Cohen, L. G. Consolidation of human skill linked to waking hippocampo-neocortical replay. Cell Rep. 35, 109193 (2021).

[60] Rasch, B. & Born, J. About sleep’s role in memory. Physiol Rev. 93, 681–766 (2013).

[61] Larkum, M. E. & Nevian, T. Synaptic clustering by dendritic signalling mechanisms. Curr. Opin. Neurobiol. 18, 321–331 (2008).

[62] Takahashi, N. et al. Locally synchronized synaptic inputs. Science 335, 353–356 (2012).

[63] Nicola, W. & Clopath, C. Supervised learning in spiking neural networks with FORCE training. Nat. Commun. 8, 2208 (2017).

[64] DePasquale, B., Churchland, M. & Abbott, L. F. Using firing-rate dynamics to train recurrent networks of spiking model neurons. *arxiv:*1601.07620 (2016).

[65] Krotov, D. & Hopfield, J. J. Unsupervised learning by competing hidden units. Proc. Natl Acad. Sci. USA 116, 7723–7731 (2019).

[66] Amato, G., Carrara, F., Falchi, F., Gennaro, C. & Lagani, G. Hebbian learning meets deep convolutional neural networks. In International Conference on Image Analysis and Processing, 324–334 (2019).

[67] Mnih, V. et al. Human-level control through deep reinforcement learning. Nature 518, 529–533 (2015).

[68] Mnih, V., et al. Asynchronous methods for deep reinforcement learning. *arXiv:*1602.01783 (2016).

[69] Such, F. P. et al. Deep neuroevolution: Genetic algorithms are a competitive alternative for training deep neural networks for reinforcement learning. arXiv:1712.06567 (2017).

[70] Miconi, T., Rawal, A., Clune, J. & Stanley, K. O. Backpropamine: training self-modifying neural networks with differentiable neuromodulated plasticity. In International Conference on Learning Representations (2020).

[71] Najarro, E. & Risi, S. Meta-learning through hebbian plasticity in random networks. In Advances in Neural Information Processing Systems (2020).

[72] Tang, Y., Nyengaard, J. R., Groot, D. M. D. & Gundersen, H. J. Total regional and global number of synapses in the human brain neocortex. Synapse 41, 258–273 (2001).

[73] Zhou, Z.-H., Yu, Y. & Qian, C. *Evolutionary Learning: Advances in Theories and Algorithms* (Springer, Switzerland, 2019).

[74] Braun, U. et al. Dynamic reconfiguration of frontal brain networks during executive cognition in humans. Proc. Natl Acad. Sci. USA 112, 11678–11683 (2015).

[75] Li, J. et al. High transition frequencies of dynamic functional connectivity states in the creative brain. Sci. Rep. 7, 46072 (2017).

[76] Hospedales, T., Antoniou, A., Micaelli, P. & Storkey, A. Meta-learning in neural networks: a survey. arxiv:2004.05439 (2020).

[77] Soysal, O. A. & Guzel, M. S. An introduction to zero-shot learning: An essential review. In 2020 International Congress on Human-Computer Interaction, Optimization and Robotic Applications (HORA) (2020).

[78] Zhuang, F., et al. A comprehensive survey on transfer learning. *arxiv:*1911.02685 (2020).

[79] Parisi, G. I., Kemker, R., Part, J. L., Kanan, C. & Wermter, S. Continual lifelong learning with neural networks: A review. *arxiv:1802.07569* (2019).

[80] Elman, J. L. Learning and development in neural networks: The importance of starting small. Cognition 48, 71–99 (1993).

[81] Sukhbaatar, S., et al. Intrinsic motivation and automatic curricula via asymmetric self-play. *arXiv:*1703.05407 (2017).

[82] Gruber, M. J., Ritchey, M., Wang, S.-F., Doss, M. K. & Ranganath, C. Post-learning hippocampal dynamics promote preferential retention of rewarding events. Neuron 89, 1110–1120 (2016).

[83] Qian, C., Bian, C. & Feng, C. Subset selection by pareto optimization with recombination. In Proceedings of the AAAI Conference on Artificial Intelligence (2020).

[84] Masse, N. Y., Grant, G. D. & Freedman, D. J. Alleviating catastrophic forgetting using contextdependent gating and synaptic stabilization. Proc. Nat Acad. Sci. USA 115, E10467–E10475 (2018).

[85] Gu, Y. et al. Abnormal dynamic functional connectivity in alzheimer’s disease. CNS Neurosci. Ther. 26, 962–971 (2020).

[86] Cheng, J. & Ji, D. Rigid firing sequences undermine spatial memory codes in a neurodegenerative mouse model. eLife 2, e00647 (2013).

[87] Iaccarino, H. F. et al. Gamma frequency entrainment attenuates amyloid load and modifies microglia. Nature 540, 230–235 (2016).

[88] Stimberg, M., Brette, R. & Goodman, D. F. M. Brian 2, an intuitive and efficient neural simulator. eLife 8, e47314 (2019).

[89] Kim, C. M. & Chow, C. C. Learning recurrent dynamics in spiking networks. elife 7, e37124 (2018).

[90] Anand, A., et al. Unsupervised state representation learning in Atari. In Conference on Neural Information Processing Systems (2019).

